# Single circuit in V1 capable of switching contexts during movement using VIP population as a switch

**DOI:** 10.1101/2020.09.24.309500

**Authors:** Doris Voina, Stefano Recanatesi, Brian Hu, Eric Shea-Brown, Stefan Mihalas

## Abstract

As animals adapt to their environments, their brains are tasked with processing stimuli in different sensory contexts. Whether these computations are context dependent or independent, they are all implemented in the same neural tissue. A crucial question is what neural architectures can respond flexibly to a range of stimulus conditions and switch between them. This is a particular case of flexible architecture that permits multiple related computations within a single circuit.

Here, we address this question in the specific case of the visual system circuitry, focusing on context integration, defined as the integration of feedforward and surround information across visual space. We show that a biologically inspired microcircuit with multiple inhibitory cell types can switch between visual processing of the static context and the moving context. In our model, the VIP population acts as the switch and modulates the visual circuit through a disinhibitory motif. Moreover, the VIP population is efficient, requiring only a relatively small number of neurons to switch contexts. This circuit eliminates noise in videos by using appropriate lateral connections for contextual spatio-temporal surround modulation, having superior denoising performance compared to circuits where only one context is learned. Our findings shed light on a minimally complex architecture that is capable of switching between two naturalistic contexts using few switching units.

**Author Summary:** The brain processes information at all times and much of that information is context-dependent. The visual system presents an important example: processing is ongoing, but the context changes dramatically when an animal is still vs. running. How is context-dependent information processing achieved? We take inspiration from recent neurophysiology studies on the role of distinct cell types in primary visual cortex (V1).We find that relatively few “switching units” — akin to the VIP neuron type in V1 in that they turn on and off in the running vs. still context and have connections to and from the main population — is sufficient to drive context dependent image processing. We demonstrate this in a model of feature integration, and in a test of image denoising. The underlying circuit architecture illustrates a concrete computational role for the multiple cell types under increasing study across the brain, and may inspire more flexible neurally inspired computing architectures.

## 1 Introduction

Our brains are unique in their ability to adapt to the context in which stimuli appear. Animals face the problem of processing visual stimuli rapidly and efficiently while adapting to different contexts every time they transition to a new environment (e.g. from jungle to savanna, from the shores of a river to underwater). A classic example of adaptation to different contexts is discussed in Barlow’s “efficient coding hypothesis” [4], which proposes that sensory systems encode maximal information about environments with different statistics [46, 47]. In this and other cases, when context changes, neural circuits switch from previous strategies of feature representation to new ones that are better adapted to the statistical properties of the new context. How the neuronal circuitry of the brain is organized to account for the multitude of contexts animals may encounter has not yet been established [62]. In particular, when do we need separate circuits for different contexts, and when can single circuits be modulated to switch among multiple contexts [23, 32, 65, 8, 38, 13, 62]? Our aim is to identify a biologically constrained network that is capable of switching contexts, and to infer the building blocks required for such switching. In constructing such a network we will only discuss and include the structural and functional detail needed for the switching of contexts.

We focus on a concrete setting in which rapid context switching is apparent. This is mouse V1, which responds differently to inputs when the animal is running (moving condition), compared to when it is stationary (static condition) [44, 20]. When the animal transitions from standing still to running, visually-evoked firing rates significantly increase. For example, in one experimental setting, the firing rate of neurons in layers II/III of area V1 more than double [44], while in layer V of V1, noise correlations between pairs of neurons are substantially reduced [15].

While an enormous diversity of cell types has been characterized [57], in this work we focus on the three primary classes of inhibitory interneurons: vasoactive intestinal peptide (VIP), somatostatin (SST), parvalbumin (PV), and one class of long range projecting excitatory neurons the pyraidal neurons (PYR) [20, 9, 53, 48] (Fig. 1a). VIP is an inhibitory population of neurons which is very strongly modulated by running [20]. In our simplified model of the circuit, VIP neurons act in a switch-like manner: they are silent when animals are static, but start firing when animals are running, inhibiting SST cells and hence releasing PYR cells from SST inhibition. The disinhibition of PYR cells is not uniform, but rather a complex pattern which is dependent on the particular PYR cell response. We will show that the switch can only be effective if PYR cells provide input information to the VIP cells. Although this simple model does not capture all the physiological responses of VIP neurons, we believe the model captures the crux of the disinhibitory switching computation at the expense of biological realism.

**Figure 1:**
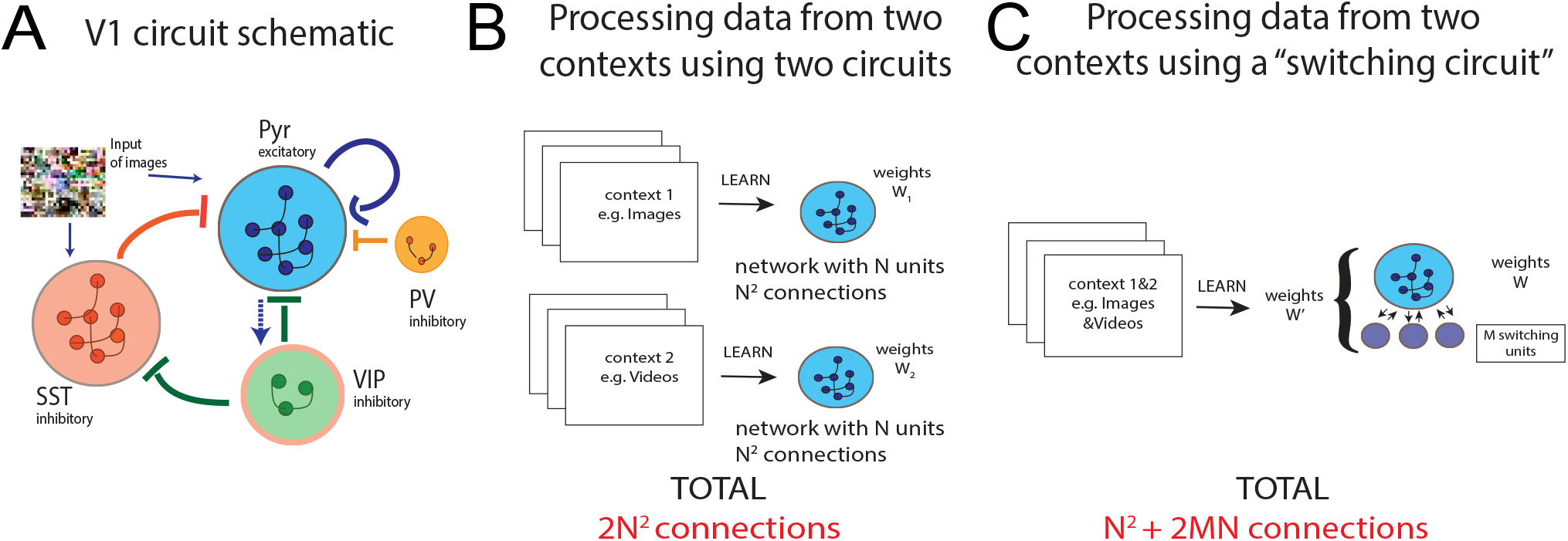
a. Schematic of circuit involving VIP, SST, PV, and PYR groups of neurons. When VIP are silent, PYR are self-excitatory, while SST and PV inhibit PYR. When VIP are active, they inhibit the PYR, while also creating a disinhibitory motif given by VIP-SST-PYR. The potential connection from PYR to VIP explored in this paper is marked with a dotted arrow. b. Processing of two input types (e.g. images, videos) happens using two separate networks for each type of input, each having *N* units with 2*N*^2^ weights in total to learn. c. Processing of two input types can be done with one circuit — a switching circuit with N units adapted to one of the contexts, and *M* switching units that turn on when the other context is presented. We may want *M* ≪ *N*, with *N*^2^ + 2*NM* connections to learn (assuming switching units are not inter-connected). When the number of switching units required in a switching circuit is small, there are fewer connections that need to be learned; more specifically, if *M* < *N* ⟹ *N*^2^ + 2*MN* < 2*N*^2^. This generalizes well to a range of circuits, including in the case of sparse connectivities, as often presented throughout the paper.

We study this circuit using a model in which the contextual information is stored in the lateral connections between neurons [26]. Each neuron receives information about the visual scene from feedforward connections (which can be arbitrary in this model), and complements this with surround information provided by nearby neurons. The connections are dependent on the statistics of the environment; more precisely they depend on the frequency of cooccurrence in the environment of the features which the neurons represent. These connections are most useful if the information from the feed-forward connections is corrupted (e.g. by an occlusions).

Importantly, the contextual information via lateral connections comes not only from the spatial surround, but also from the past. Synaptic delays introduce a constraint on the available information each neuron gets. During the static condition, past surround information matches present information, and thus there is no temporal variability of the context. During movement, this no longer holds; neighboring features now also vary temporally, which changes the cooccurrence frequency, and hence the statistics of the moving context is different. We aim to find connection strengths from the switching VIP units that, during movement, modulate firing rates and neuronal correlation structure to adapt and enhance encoding of visual stimuli when the moving context is turned on. Although throughout the paper we focus on the visual circuit and the switching role of the VIP neural population, these results can be generalized to circuits processing multiple contexts, and thus their applicability has broader scope. In the discussion section, we list several other biological examples of circuits processing multiple contexts.

Understanding switching circuits may also further aid efforts to design both flexible and efficient artificial neural architectures. This research area has benefited from bio-inspired architectures and algorithms like elastic weight consolidation [30], intelligent synapses [64], iterative pruning [37], leveraging prior knowledge through lateral connections [54], task-based hard attention mechanism [52], block-modular architecture [58], etc. to enable sequential learning by eliminating “catastrophic forgetting” (where previously acquired memories are overwritten once new tasks are learned). We hypothesize that a few switching units akin to VIP can be incorporated as part of the hidden layers to enable context modulation. This makes such a switching circuit architecture (Fig. 1c) more efficient than employing separate circuits for the different contexts (Fig. 1b) because switching circuits have fewer connections to learn ^1^. We hope such a circuit architecture will inspire next-generation flexible artificial nets that can process stimuli in changing contexts.

### Outline of paper

In section 2.1, we first detail a model introduced in [26] that describes neuronal connections and firing rates of a circuit adapted to static visual scenes (images). We next extend this model to the case of circuits adapted to moving visual scenes (videos). These circuits are attuned to the statistical regularities of movement and take into account constraints of biological networks, like synaptic delay. We are able to map these two circuit models to the V1 circuit, consisting of PYR, SST, and PV neuron populations. We thus obtain two different networks with full cell-type specifications achieving optimal context integration for static and moving contexts, respectively. In section 2.2 we detail the datasets and procedures used to quantify connectivities and firing rates in these two circuits. In section 2.3, we go on to describe a circuit that can switch between neuronal activity in static circuit and neuronal activity in the moving circuit, by virtue of adding a single population, the VIP. We find that VIP projections to SST and PYR are not enough to shift activity during movement, but that we need a feedback connection from the PYR to the VIP (section 2.4). The resulting circuit is the minimally complex circuit resembling V1 we have found to switch contexts. In section 2.5, we describe how this circuit switches using only a small number of VIP units. We follow up on these results in section 2.6, where we utilize this switching circuit to obtain better reconstructions of videos in conditions of high noise. Finally, we evaluate the new switching circuit architecture with data from V1 that confirms some of the model’s predictions (section 2.7).

## 2 Results

### 2.1 Theoretical models of processing visual information in static and moving contexts

We introduce a model of visual processing where feedforward and lateral connections between neurons serve different roles. The lateral connections between neurons perform unsupervised learning of the probability of co-occurrence of features in the visual space which the neurons represent. For the purpose of this study, the feedforward connections can be arbitrary, and the microcircuit described here can be at any level of processing. This separation of the roles for the feedforward and lateral connections allows for an easy implementation of both supervised and unsupervised learning in deep networks [27].

Here, we show how this model can integrate information from the surround using these within-layer connectivities in both static and moving states. However, integration of these two contexts results in two distinct circuits needed to perform visual processing under different conditions (static vs moving). The model optimally integrates context in the Bayes sense, meaning it uses priors on the co-occurrence of features in natural scenes when integrating information from the surround. These priors reflect the known statistical regularities of the environment [55, 4, 39] and weigh the surround contributions appropriately. We are then able to map this model formalism to the circuit architecture in V1 described above while specifying steady state network weights and activations, as well as cell type functionality. This model emphasises robust coding, and applies best in conditions of high noise, where parts of the visual scene are missing due to occlusions or are corrupted, and thus where context information may play a critical role. We next describe our model of visual processing in detail.

#### Model of visual processing in the static context

To study optimal context integration in the static condition (where the visual input is static images), we take as a starting point a model proposed by Iyer et al. in [26] where model neurons respond to a patch in the visual space — the classical receptive field — but this response is modulated by a larger region of space — the extra-classical receptive field. The extra-classical receptive field contribution is determined by nearby local receptive fields providing indirect input from a larger area of visual space (Fig. 2a). Specifically, inter-neuron interactions providing extra-classical information from the surround via lateral connections (cfr. Methods sec. 4.1) complement intrinsic neuronal responses to classical receptive fields to determine firing rates.

**Figure 2:**
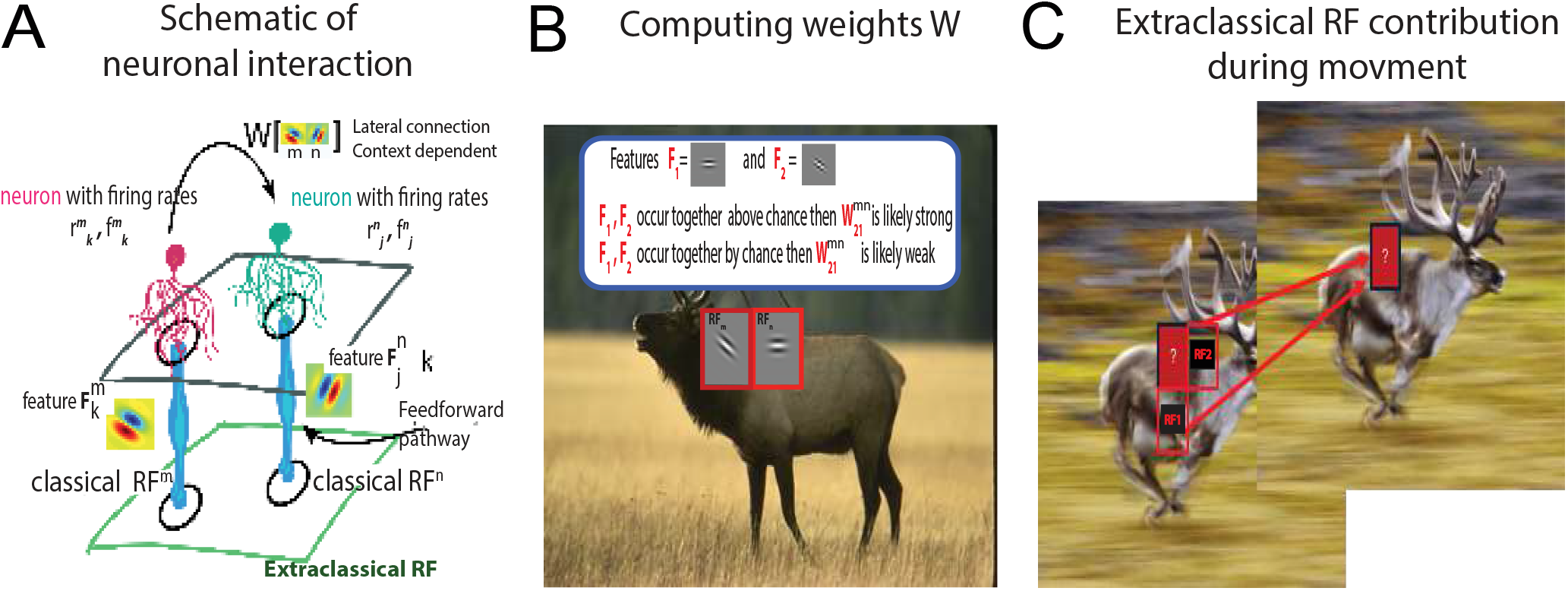
a. Neurons receive stimulus input from a patch in space at position *n*, their classical receptive field (RF), but also from surrounding patches in space (for e.g. the patch at position *m*) through interactions with other neurons. These neurons are connected by weights 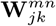 that depend on the statistical regularities of natural scenes. b. When features **F**_1_ and **F**_2_ at positions m,n occur together often in natural scenes, then 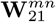 is strong; when **F**_1_ and **F**_2_ occur together by chance, without significant correlation, 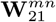 is close to 0. c. Spatio-temporal surround for motion processing. Due to synaptic delay, context integration uses surrounding patches that are also Δ*t* ms in the past to assess the features in the present frame.

Starting from the assumption that firing rates of a population of neurons encode the probability of specific features being present in a given location of the image, we consider a probabilistic framework that includes probability of feature occurrence and feature co-occurrence, that we can then map to an equation involving firing rates of neurons and weights (cfr. Methods sec. 4.1). In general, a feature *j*, denoted by **F**_*j*_, describes a specific pattern that neurons are most attuned to, that can vary from simplistic, like Gabor filters, to complex, like faces or objects that are robust to stimulus transformations such as scale and position changes. In more detail, for neurons responding to 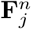 (feature *j* at patch *n*), we define 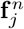 to be the steady-state firing rate due to the classical receptive field, and 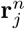 to be the (overall) steady-state firing rate taking into account the extra-classical receptive field contribution. The probabilistic assumption stated above is such that 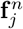 relates to the probability 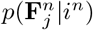 by the following relation:

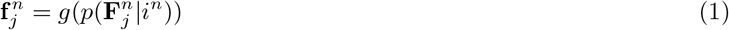

where *g* is a monotonically increasing function, *i^n^* is a patch *n* in visual space, and 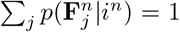. For simplicity, we fix *g* to be the identity, leaving the relaxation of this linear assumption for future work. With 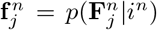, neurons tuned for distinct features respond differently to the same patch *i^n^* in visual space depending on how well its corresponding feature is represented. Operationally, to compute 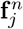 in response to an image, we first chose a basis of features, for e.g. features obtained by approximating spatial receptive fields from recorded neurons in V1. We then pre-processed the image (cfr. Methods 4.2), convolved the image with feature *j* and normalized the result such that the sum over all features is 1 at each spatial position, and finally considered the patch *i^n^* of the normalized convolution.

Once 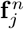 is computed, we can continue assuming that neuronal firing rates contain information about feature occurrence in the surround, so that 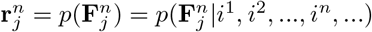, then using Bayes rule to express this in terms of feature probability at patch *i^n^* and at surrounding locations, and finally mapping the resulting equations to neurobiological quantities. These operations yield that the firing rates 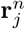 of neurons are the result of modulating the classical receptive field firing rate 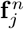 by extra-classical receptive field information from the surround which is a linear function of other neurons’ classical receptive field firing rates, 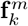. These firing rates are weighed by the lateral connections **W**^static^, representing the prior information about the statistical regularities of natural images. After ignoring terms which are due to higher order modulation of the surround (cfr. Methods sec. 4.1), specifically neurons from the surround having surround modulation of their own, we obtain the following firing rates as exemplified in the schematic in Fig. 2a and explained in detail in Methods sec. 4.1:

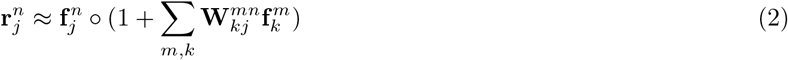

with the weights expressed as:

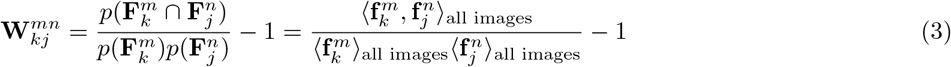

where 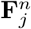 is a Gabor-like feature *n* at location *j* that we will illustrate shortly, the symbol ⋂ denotes the co-occurrence of two features, and ○ is the Hadamard product, the element-wise multiplication between tensors 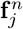 and 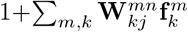. Further, 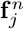 is the evoked firing rate due to the classical receptive field of neurons firing for feature 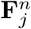, and 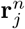 is the firing rate of neurons firing for feature 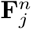 using information from classical and extra-classical receptive fields. The sum 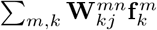 is over neurons with receptive fields at different locations *m*, responsive to features *k*. Finally, 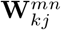 is the connectivity in the static context between neurons responsive to features 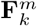 and 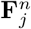. We define 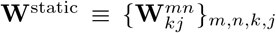 as the connectivity applied to static visual scenes. Assuming that weights only connect neurons with non-overlapping receptive fields, the resulting weights are sparse (see Methods sec. 4.2).

From a computational perspective, the organism cannot measure the feature probabilities and joint probabilities in (1) and (3) directly, but these can be estimated given our defined neural code as the convolutions between image and feature, i.e. 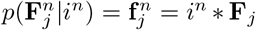, and as the cross-correlations between classical receptive field firing rates, i.e. *p*(**F**_*k*_ ⋂ **F**_*j*_) = 〈**f**_*k*_, **f**_*j*_〉. By mapping these probabilistic statements on feature occurrence to neurobiological quantities that capture firing rates and weights, we have obtained a circuit that does approximate context integration, extracting information through priors embedded in the neural connectivities. While the start of the model is Bayes-optimal via Equations (36) – (38), a set of approximations are needed to keep the circuit simple.

There are multiple possible mappings from the probabilistic framework to the neurobiological circuit [26], but the current correspondence is straightforward and yields successful predictions from data, such as like-to like connectivity, as detailed below. When a pair of features is frequently co-occurring, weights between neurons preferential for these features are strong and positive (Fig. 2b). In contrast, when two features are unlikely to co-occur in the same image the connectivity is strong and negative. Overall occurrence probabilities of individual features normalize the co-occurrence probabilities so that the weights express the co-occurrence of features over and above chance. Co-occurrence probabilities of features are then averaged over many natural scenes so that the corresponding weights **W**^static^ capture the statistical regularities of natural environments.

#### Model of visual processing in the moving context

We next show how the framework above can be applied to the moving context. While Equations (2) – (3) show how connectivity and firing rates can be optimized to account for spatially co-occurring features — features that appear at the same moment in time but in different locations of the visual field — we now extend these equations to account for temporal co-occurring features — features which occur at nearby moments in time at different locations of the visual field.

In more detail, context is generally integrated from Δ*t* in the past due to synaptic delay (Fig. 2c), and weights are proportional to co-occurrence probabilities of neighboring features that are also separated by a time window Δ*t*. This is a direct generalization of the model in [26] to the time domain, and includes synaptic delay as a biologically motivated constraint. The extended model can capture how local circuit connectivity is shaped by spatio-temporal correlations across receptive fields and across time windows characteristic of biological processes like synaptic delay. The firing rate during the moving context is (cfr. Methods sec. 4.2):

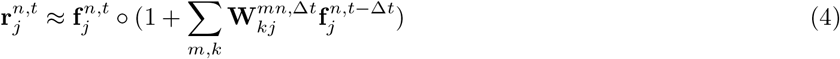

with the weights expressed as:

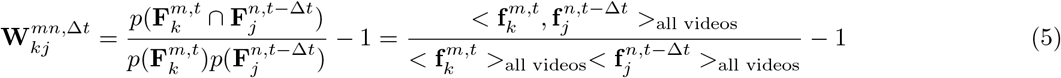

where we apply an analogous notation as for Eq. (2) and Eq. (3), the only difference being the additional *t*, Δ*t, t* − Δ*t* superscripts that denote the time coordinate for the features, firing rates, and weights. 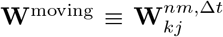 is the connectivity in the moving context between neurons responsive to features 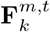 and 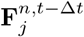 whose activation is separated by a time delay Δ*t*. Note that the expression for 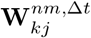 as shown in (5) also holds for the static context when we use static visual input to compute the weights, such that **f**^*t*^ = **f**^*t*-Δ*t*^, for all *t*, Δ*t*.

### 2.2 Modeling firing rates and weights in networks responding to images and videos

We next describe two separate circuits capable of doing optimal context integration in each of the moving and static contexts. We characterize these two circuits through the connectivities **W**^static^ and **W**^moving^, computed by using images and videos in training datasets and applying formulas (3) and (5). Once the corresponding connectivities are specified, we can further characterize the static and moving circuits by their neural activations. In the following, we elaborate, section by section, on the algorithm we implemented to compute the static and the moving weights.

#### Dataset and feature preparation

We applied our framework for processing static images and videos to different benchmark datasets, chosen to address differences in the statistics of visual features across conditions: during viewing of static images (static condition) and during viewing of videos which contain motion (moving condition). For the static condition, we used 300 selected grayscale images of the BSDS dataset [40] (Fig. 3a) while for videos, the BSDS dataset is pre-processed through a smaller sliding window that travels along the image to reproduce motion (Fig. 2b, cfr. Methods sec. 4.4). Although in general the sliding window can move in any direction (see Figs. S1 and S2 for results in this case), here we constrained it to move solely in the horizontal direction to roughly approximate flow of images across the (sideways-facing) eyes of mice during forward movement. We have not used a generic dataset of natural videos since most videos in such datasets contain limited movement of objects, humans, or animals, rather than movement of sections of an environment that would mimic the visual experience of a running animal.

**Figure 3:**
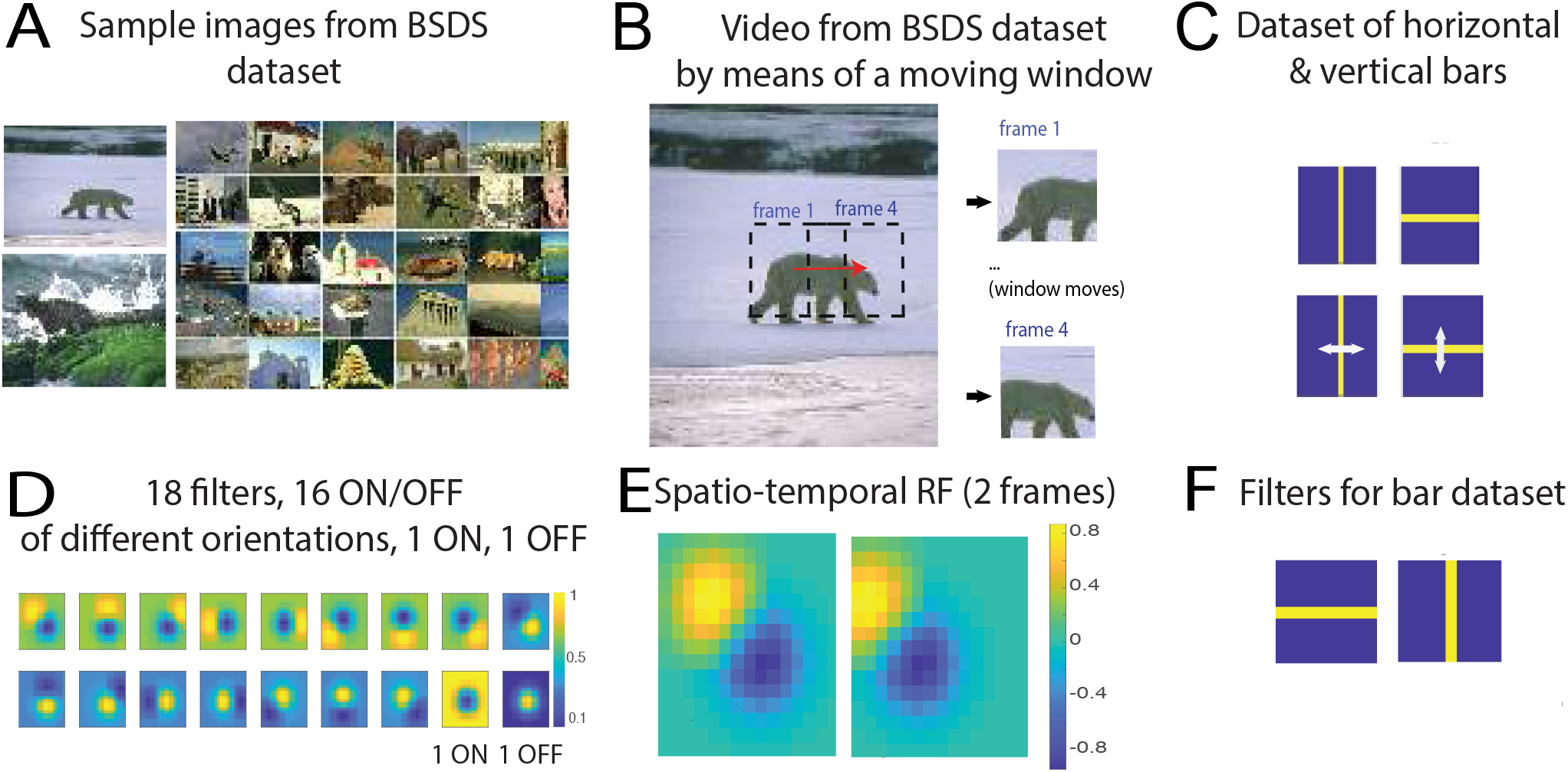
a. Sample images from the BSDS dataset. Images of animals, human faces, landscapes, buildings, etc. are used. b. Sliding window on images from the BSDS dataset so that the appearance of movement is achieved. Shown by the red arrow is how much the window has moved from frame 1 to frame 4. In general, movement of sliding window is random and in any direction, but we focus on horizontal movement in the case of natural videos. c. Images of horizontal and vertical bars (above) and how the bars move in videos (below). d. 18 filters: ON, OFF, ON/OFF with 2 Gaussian subfields, different subfields dominating, at different intensities and orientations. Colorbars show the different intensities of pixels. e. Example of a spatio-temporal filter comprising of 2 frames. Spatio-temporal filters are added to the 18 original filters, to make up a total of 34 filters. The filter shown here over 2 frames captures a 45 deg bar moving to the left and is obtained by translating the original filter by 3 pixels. Colorbars show the different intensities of pixels to the left. f. 2 filters for the simplistic “bar world” comprising of a horizontal and a vertical bar, respectively.

We generated a dictionary of features (filters) based on a parametrized set of models derived from recordings in V1 [19]. This contains 18 filters with Gaussian subfields (Fig. 3d) at different relative intensities and orientations. We added filters containing a temporal dimension — *spatio-temporal filters* — to obtain a set of 34 filters. Our spatio-temporal filters consist of 2 frames (Fig. 3e) and represent a temporal shift by several pixels in the horizontal direction, corresponding to the direction of movement and amount of displacement of the sliding window in the videos described above.

To more easily illustrate and interpret our model, we first tested our framework on a different, synthetic context. We analyzed a simplified 9 × 9 world of horizontal and vertical bars moving up-and-down as well as left-and-right (Fig. 3c). This simple dataset has only two features, horizontal bars and vertical bars (Fig. 3f), but movement can be in any of the four orthogonal directions.

#### Computing the weights W^static^, W^moving^

The firing rates **f** due to the classical receptive field represent feature probabilities (Equation (1) with *g*(*x*) = *x*) and were computed by the following sequence of operations: pre-processing inputs and filters (cfr. Methods sec. 4.2), convolving the image or video frames with the respective sets of filters, rectifying, and then normalizing so that all firing rates 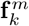 lie in the interval between 0 and 1 and sum up to 1 across all features *k*. To find the weights for static and moving contexts, **W**^static^ and **W**^moving^, we fixed Δ*t*. After convolving 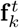 and 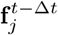 in accordance with Equations (3), (5), and following the procedure outlined in Fig. 4a, we obtained a high dimensional tensor that characterizes the connections between each pair of cell types (*k, j*) at each position in the image. Using the feature 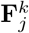 as a proxy for an excitatory cell “type,” the resulting tensor is 4 dimensional, with dimensions: cell type of the source, cell type of the target, and relative spatial position of the source and target in *x* and *y* directions.

**Figure 4:**
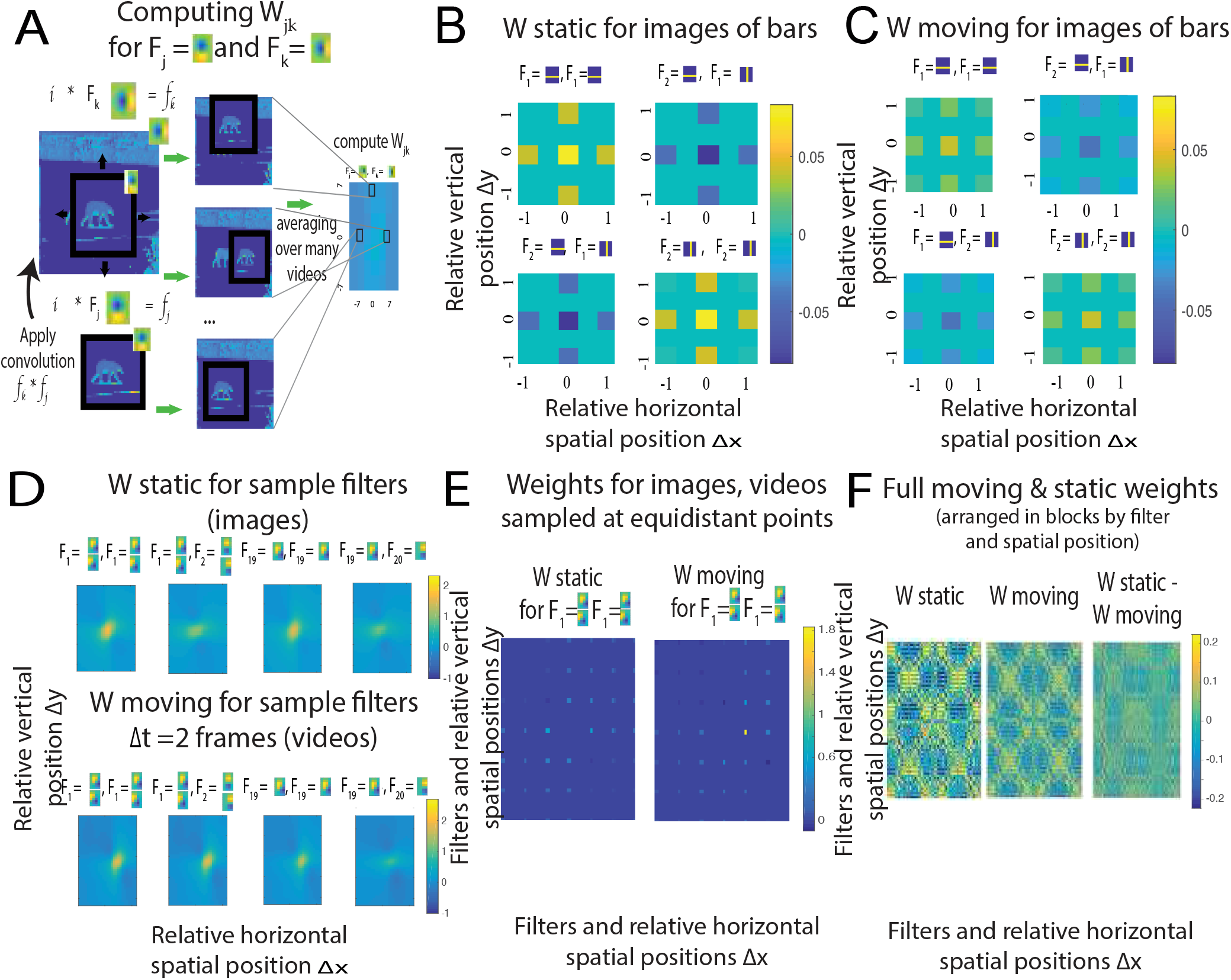
a. Schematic of how weights are represented. Instead of representing weights corresponding to all pairs of patches in visual space, we assume neighboring patches elicit the same connectivity regardless of where in the visual field the receptive field is (weights obey the property of translational invariance). b. Static weights for the dataset of images of bars. c. Moving weights for the dataset of videos of bars. d. Static weights (up) and moving weights (down) for the dataset of natural images/videos during horizontal motion only. e. Sparse versions of slices from the static and moving weights for the datasets of natural images/videos during horizontal motion. Weights between neurons whose receptive fields are not at certain pre-selected, sufficiently far apart locations in the visual space were discarded to satisfy the constraint that patches are independent. f. The full (non-sparse) tensors **W**^static^, **W**^moving^, and **W**^moving^ – **W**^static^, ordered first by spatial position, then by filter.

#### Simplifications to weights

We make three simplifications to reduce the number of parameters in this tensor (cfr. Methods 4.2): (1) we assume translational invariance so that only the relative position of two filters is relevant (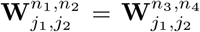 when 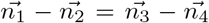); (2) the model is designed to compute connections to neurons which receive independent observations, thus we only consider connections between neurons whose receptive fields are sufficiently far apart (i.e. at least half a receptive field apart), (3) as statistical dependencies in natural images decay with distance, we limit the spatial extent of connectivity to three times the size of the classical receptive field. Fig. 4b shows several 2D slices through this tensor, corresponding to a specific cell source and target, as well as the full static and moving weights (figs. 4b to 4f) ordered by spatial position and feature type (see also Fig. S1). Figures 4b and 4c serve to provide some intuition as to what these weights represent and how they are structured: in the dataset of bars, horizontal feature **F**_1_ frequently occurs or is absent together with other horizontal features **F**_1_ at neighboring locations, which leads 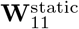 to have positive values. Conversely, horizontal feature **F**_1_ occurs always when vertical feature **F**_2_ is absent, and viceversa, leading to negative weights 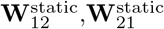 (Fig. 4b).

#### Characterizing W^moving^ in the case of two different video statistics

In the generation of the video dataset we use a sliding window to enforce controlled and comparable statistics between the moving and static contexts. When the sliding window is free to move in all directions, the moving weights tend to be weaker in absolute value, which holds for the simple dataset of bars (figs. 4b to 4c), and the weights generated from the dataset of natural images and videos (figs. S1a to S1b). This effect is due to the weaker statistical dependence of features separated by the time window Δ*t*. Feature co-occurrence, and thus connectivity, is affected by the distortions during movement, like change of orientation of objects, or appearance or disappearance of objects in the visual scene. Moving weights in this case are approximately a smoothed out versions of the static weights (figs. 4b to 4c, figs. S1a to S1b). In these conditions, as the information from surround is less reliable, the feedforward input plays a more important role during movement.

In the case when the sliding window moves *s* pixels horizontally in Δ*t* time steps, 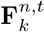 and 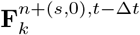 actually coincide so that their probability of co-occurrence is maximized. This means that for horizontal movement, 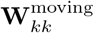 peaks *s* pixels from the center for any feature **F**_*k*_ and 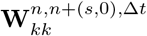 is strong (figs. 4d to 4e). Results for natural videos below are for horizontal movement, although the same general conclusions hold when movement is allowed in any direction (see Fig. S3).

Finally, using **W**^static^, **W**^moving^ and applying Equations (2), (4), we obtain the corresponding firing rates **r** in both static and moving contexts.

### 2.3 Implementing a switching circuit

Having two just defined the two optimal connectivities, **W**^static^ and **W**^moving^, for the static and moving contexts, we next consider whether a single circuit involving the cell types described above (VIP, PYR, SST, and PV) can respond optimally in these two contexts and switch between them. We additionally seek the computational principles behind the minimally complex circuit (i.e. the circuit with fewest connections) for such a switching circuit. Specifically, we ask whether a circuit with optimal weights for the static context can switch to produce nearly optimal activities in the moving context, via projections from a set of switching units. In such a circuit every PYR neuron approximates Bayesian inference, combining classical receptive field information with information from the surround to estimate feature probability.

We start by rewriting the model described by Equations (3) – (4) in vector form to obtain the following firing rates:

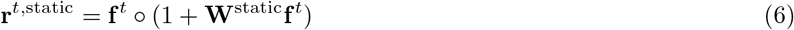

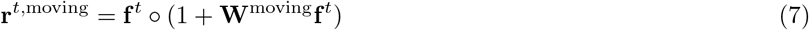

Assuming, as discussed above, that the activation of the VIP neural population implements the switch between contexts, we want the switching circuit to reproduce the firing rates given by (6) when the VIP neurons are silent in the static context, and the firing rates given by (7) when the VIP neurons are active in the moving context (Fig. 5a). We next explain how **r**^static^, **r**^moving^ above can be modeled as the firing rates of the PYR neurons.

**Figure 5:**
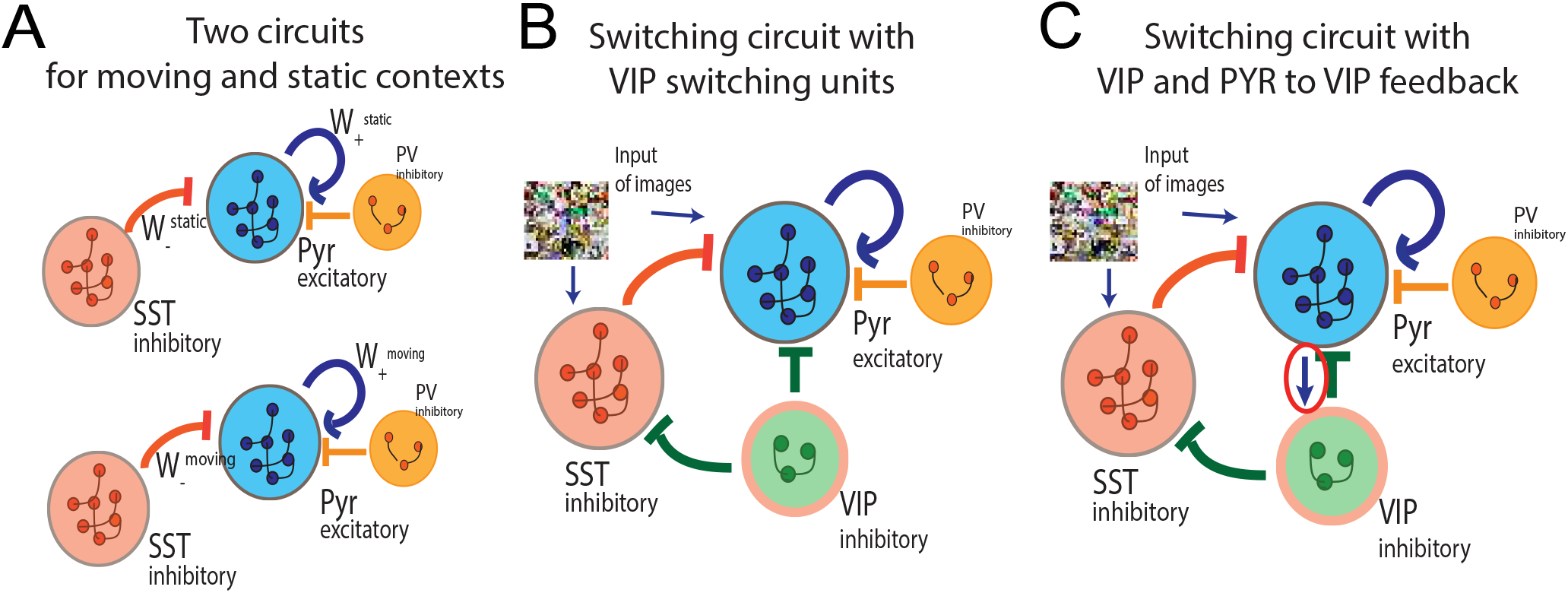
a. Two separate circuits for optimal visual processing of static (top) and moving contexts (bottom), respectively. b. The proposed switching circuit with the VIP population approximates the static circuit when the VIP are silent and the animal is static, and approximates the moving circuit when the VIP are active and the animal is moving. c. Previous circuit, but with a feedback connection added from the PYR population to the VIP.

When the VIP are silent, the only groups of neurons active are PV, SST, and PYR. This circuit is equivalent to one without any VIP connections, reproducing firing rates of PYR given by (6) when the animal is static. PYR neurons contribute to integrating surround information through excitatory projections, and receive inhibitory feedback from SST interneurons [7]. PV implements a normalization of the PYR population in our model, consistent with data on their connectivity [28, 48]. Empirically it has been shown these neurons receive the average inputs of the PYR neurons whose receptive fields overlap with their classical receptive fields, and project back equally [48]. In our model, this normalization applies to the classical receptive field **f**, as described in Methods sec. 4.2. As for the role of PYR and SST, given that PYR are excitatory and SST are inhibitory, and that 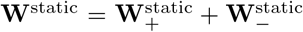, it is natural to map the positive component of the static weights, 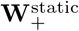, to the connections within the PYR population, and the negative component of the static weights, 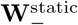 to the inhibitory connections from SST to PYR. Hence, we obtain the following:

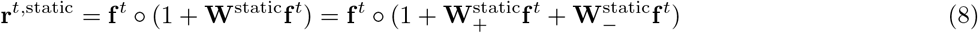

can be mapped to

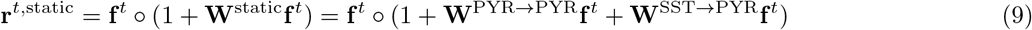

where **W**^*X→Y*^ denotes the weights that connect neuronal populations **X** (the source) and **Y** (the target).

On the other hand when VIP are active, PYR firing rates ought to reproduce the activity given by (7). We make the simplifying assumptions that the switch from static to moving can happen instantaneously, and that the VIP switch is binary. When the animal initiates movement and the VIP turns on, the model circuit should approximate the optimal response of PYR neurons resulting from the **W**^moving^ connectivities, within a circuit where the 4 neuronal populations interact (Fig. 5b). For VIP modulation of PYR (which is either direct or through the SST) that gives rise to the optimal firing rates in the moving context, we have:

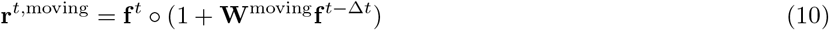

is mapped to

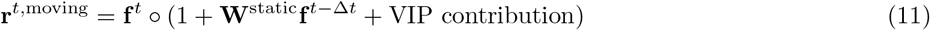

Thus, the switch in the circuit occurs as VIP neurons modulate SST and PYR neurons and make PYR switch firing rates from **r**^static^ to **r**^moving^. We now proceed to find the unknown connectivities, from VIP to PYR and from VIP to SST, that causes this to occur within the circuit (Figs. 5b to 5c).

### 2.4 In the absence of feedback to VIP neurons, the circuit is unable to switch from static to moving conditions

We attempt to describe the computational principles of the minimal switching circuit inspired by the V1 circuitry whose main structure and logic was described in [20]. After adding the switching population VIP, the goal is to find connectivities from VIP to the other two neuronal populations (PYR, SST) that would account for the PYR firing rates that yield optimal representation in the moving context. With the VIP contribution, the firing rate of PYR neurons can be expressed as (cfr. Methods sec. 4.5):

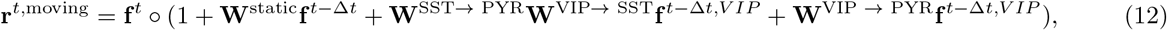

where **f**^*t*^, **f**^*t*−Δ*t*^ are firing rates due to the classical receptive field at times *t* and *t* − Δ*t* and inferred from the dataset of natural videos as outlined in sec. 2.1 and Methods sec. 4.2, **f**^t,*VIP*^ are the intrinsic firing rates of the VIP at time *t*, and **r**^*t*,moving^ is the firing rate during the moving context with the extra-classical receptive field contribution. Here, **W**^SST→ PYR^ are weights from SST to PYR, **W**^VIP→ SST^ are weights from VIP to SST, and **W**^VIP → PYR^ are weights from VIP to PYR. VIP neurons project to PYR neurons directly via weights **W**^*VIP→PYR*^ and indirectly via the SST population. The effects of the indirect pathway VIP-SST-PYR can be captured by taking the product of connectivities, yielding **W**^*SST→PYR*^**W**^*VIP→SST*^. The three unknown variables are then **f**^t,VIP^, **W**^VIP→ SST^, and **W**^VIP → PYR^, but since we assume **f**^*t, VIP*^ is constant in time *t*, this tensor can be combined with the connectivities to form the effective parameters **w**^*α*^ = **W**^*VIP→SST*^**f**^*VIP*^ and **w**^*β*^ = **W**^*VIP→PYR*^**f**^*VIP*^ and hence reduce the number of unknowns and simplify notation. Our objective is to have firing rates in the switching circuit be as closely matched as possible to the firing rates in the separate moving circuit with **W**^moving^:

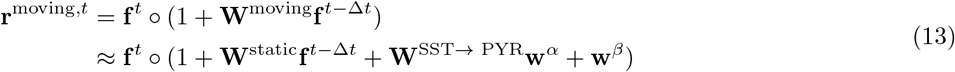

This amounts to minimizing the loss function defined by the approximation error *E*_*switch*, 1_ over the variables **w**^*α*^, **w**^*β*^:

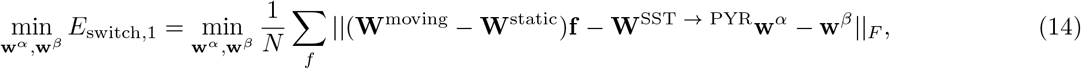

where ║·║_*F*_ is the Frobenius norm of a tensor, for all **f** (firing rates due to classical receptive fields) corresponding to video frames, and *N* is a normalization factor, the number of video frames in our dataset. **f** is inferred through our model from the datasets of video frames and features using 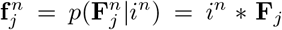 and thus is a known quantity throughout the optimization. Importantly, since **f**^*VIP*^ are firing rates and hence **f**^*VIP*^ ≥ 0, while **W**^SST→ PYR^ ≤ 0, **W**^VIP→ SST^ ≤ 0, and **W**^VIP→ PYR^ ≤ 0, we have that **w**^*α*^, **w**^*β*^ ≤ 0, and **W**^SST→ PYR^**w**^*α*^ ≥ 0.

This is a high dimensional constrained optimization problem with the loss function defined as in (14), which we solved by means of a gradient descent method using the gradient-based Adam optimizer, implemented in pytorch^2^. The weights **w**^*α*^ and **w**^*β*^ are unknown and learned by Stochastic Gradient Descent (SGD), while **W**^moving^, **W**^static^, **W**^*SST→PYR*^ ≡ [**W**^static^]_−_ are fixed. Finding the global minimum of the loss function is difficult, but the main goal is to find weights that give a small enough error *E*_*switch*, 1_ instead and later test these on a specific task to demonstrate that the optimal moving circuit can be approximated successfully (Section 2.6). We assessed the stability of our optimization by modifying several learning hyperparameters: learning rate (ranging from 0.001 to 0.1), optimization algorithm (SGD, AdaGrad, RMSProp, Adam), etc. and checking the generalization error on a small number of frames (50) that were not used during training.

Regardless of hyperparameters, our optimization procedure did not find weights that together approximate the moving circuit significantly better than the static circuit. In other words, adding VIP neurons in an attempt to switch contexts does not lead to a significantly better approximation of the moving circuit than having no VIPs. This result holds for both the simple dataset of horizontal and vertical bars, and for the more complex dataset of natural images and videos (figs. 6b to 6c).

**Figure 6:**
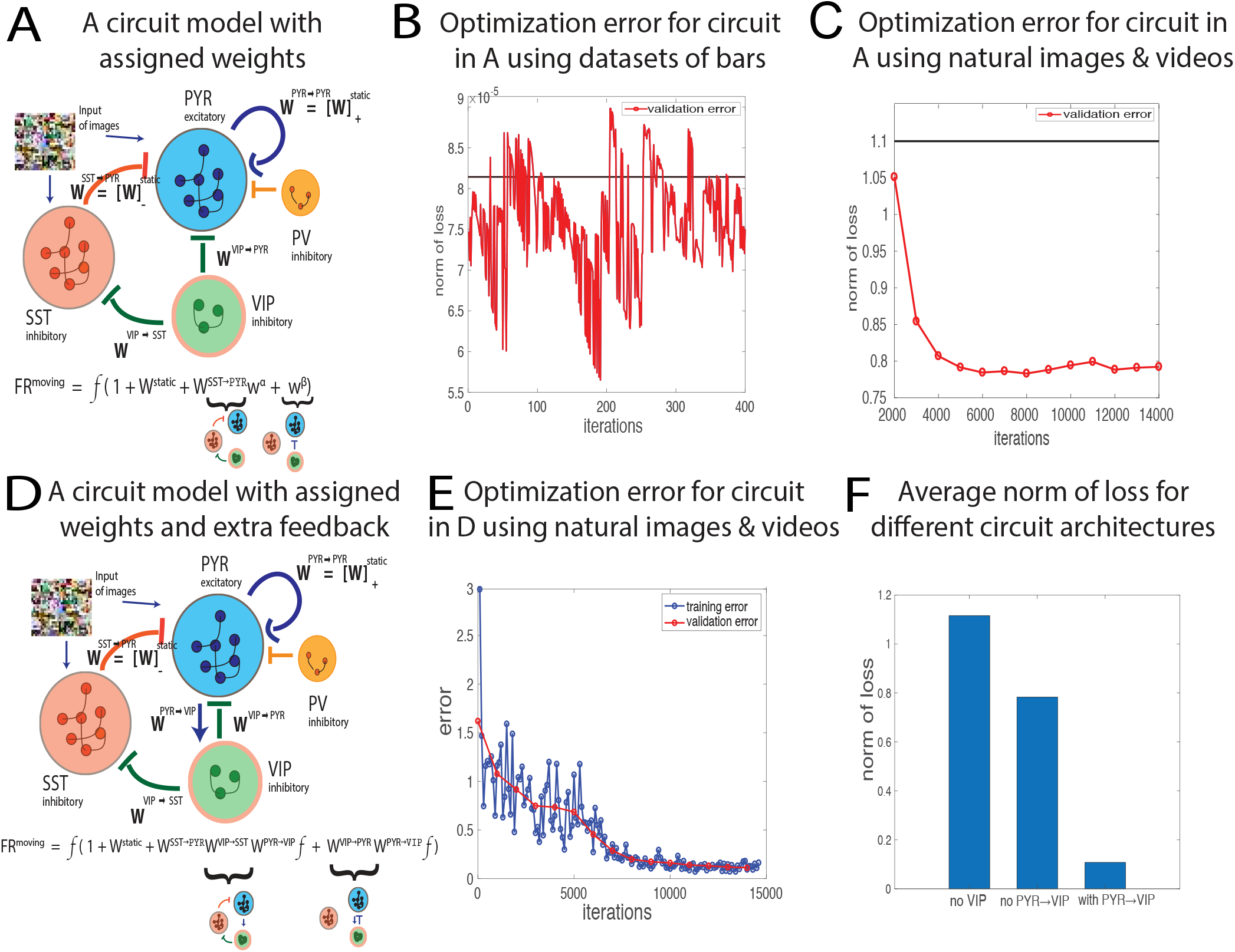
a. Goal: instead of two separate circuits for visual processing of static and moving contexts, the proposed circuit approximates the static circuit when the VIP are silent and the animal is static, and the moving circuit when the VIP are active and the animal is moving. b. Generalization/validation error found during the optimization to minimize the functional *E*_*switch*,1_ for the datasets of static and moving bars does not converge. c. Generalization/validation error found during the optimization to minimize the functional *E*_*switch*,1_ for the datasets of natural images and videos converges, but the norm of the loss function decreases by only ≈ 25%. d. Circuit as in (a), but with a feedback connection added from the PYR population to the VIP. e. Training error (blue) and generalization/validation error (red) found during the optimization to minimize the functional *E*_*switch*,2_ (movement approximation error) for the datasets of natural images and videos converges to yield a relatively small error. f. The movement approximation error for various circuit architectures: the static circuit with no VIP switching units, the circuit depicted in (a) without PYR to VIP feedback, the circuit depicted in (d).

In order to understand the origin of this failure, we mathematically analyzed the circuit at hand. Analytically, if the loss is small E_switch,1_ ≈ 0, then (**W**^moving^ — **W**^static^)**f** ≈ **W**^SST→ PYR^**w**^*α*^ + **w**^*β*^, where **f** is unique to each image in the data. The left hand side becomes a term that varies across a wide range of video frames, while the right hand side is a constant term incorporating the weights we are solving for: **w**^*α*^, **w**^*β*^. This suggests that the failure of our optimization procedure to yield weights that approximate the moving circuit results from the VIP having no stimulus dependence.

We conclude that the circuit switching between static and moving contexts must be more complex than the simple circuit here, which has only outgoing projections from VIP. Below, we introduce recurrent connections which make the VIP input-dependent, and overcome the limitations above.

### 2.5 VIP circuit with feedback from the PYR cells can switch context integration from static to moving conditions

Above we showed that a minimal switching circuit with only outgoing projections from the VIP units is insufficient to switch between the two contexts. Hence, we added an additional connection between PYR and VIP, such that the VIP group of neurons has access to information about the visual input through PYR (Fig. 5c). In this case we can approximate the firing rate of PYR during movement as follows, using the same conventions and assumptions as before (cfr. Methods sec. 4.5):

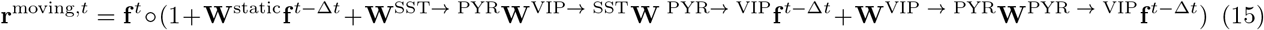

We remind the reader that **f** is the contribution to the firing rate of the classical receptive field, **W**^X→Y^ are the weights from population X of neurons to population Y of neurons, where X, Y are the PYR, SST, VIP neurons. In addition to the fixed **W**^static^ and **W**^moving^, we also fix **W**^SST → PYR^ = [**W**^static^]_−_. A schematic of the underlying circuit model, along with the corresponding formula for the firing rate of PYR, is shown in Fig. 6d.

We would like to find the three unknown weights **W**^*VIP→PYR*^, **W**^*VIP→SST*^, and **W**^*PYR→VIP*^, to best achieve the approximation:

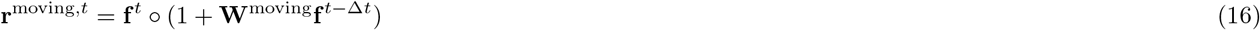

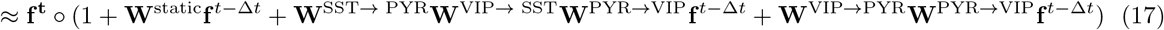

We denote the approximated expression of (17) by **r**^approx^. This approximation **r**^approx^ ≈ **r**^moving^ amounts to minimizing the loss function defining the *movement approximation error E*_*switch*,2_:

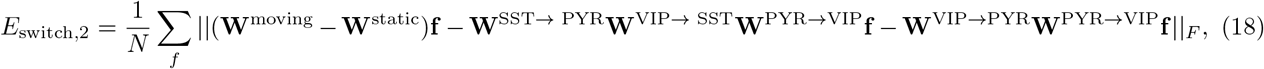

for all *N* frames whose corresponding classical receptive field firing rate is **f**. In the case of simple images and videos of bars we consider **W · f** to be the regular matrix vector multiplication, while in the case of natural scenes we perform the convolution operation **W ∗ f**. Applying convolution for natural images and videos fits with the assumption we have applied for the PYR, SST populations, that weights between neurons are translationally invariant, and further reduces the number of parameters.

To solve this high dimensional optimization problem, we set up, as in Sec. 2.4, an optimization problem with the loss function being the Frobenius norm as defined in (18). Weights to and from VIP are unknown (**W**^VIP→SST^, **W**^VIP→PYR^, and **W**^PYR→VIP^) and learned by SGD, while **W**^moving^ − **W**^static^, **W**^SST→ PYR^ are fixed. Importantly, Dale’s law is enforced (**W**^VIP→SST^, **W**^VIP→PYR^ ≤ 0, **W**^PYR→VIP^ ≥ 0) for biological realism.

To find how many switching units are needed, we varied the number of VIP neurons, which was equivalent to varying the dimensionality of tensors **W**^VIP→SST^, **W**^VIP→PYR^, and **W**^PYR→VIP^. We found the smallest number of switching neurons VIP that enabled the loss (18) to be minimized. First, for an image/video set which was 9 × 9 with horizontal and vertical bars, the loss was minimized with at least 20 VIP neurons (Fig. 7a). For comparison, there are 162 PYR and SST neurons, one for each filter and pixel in the image or frame. As increasing the number of VIP units further does not decrease the loss function, so we conclude that, for the case of barlike images, having 20 switching units is enough.

**Figure 7:**
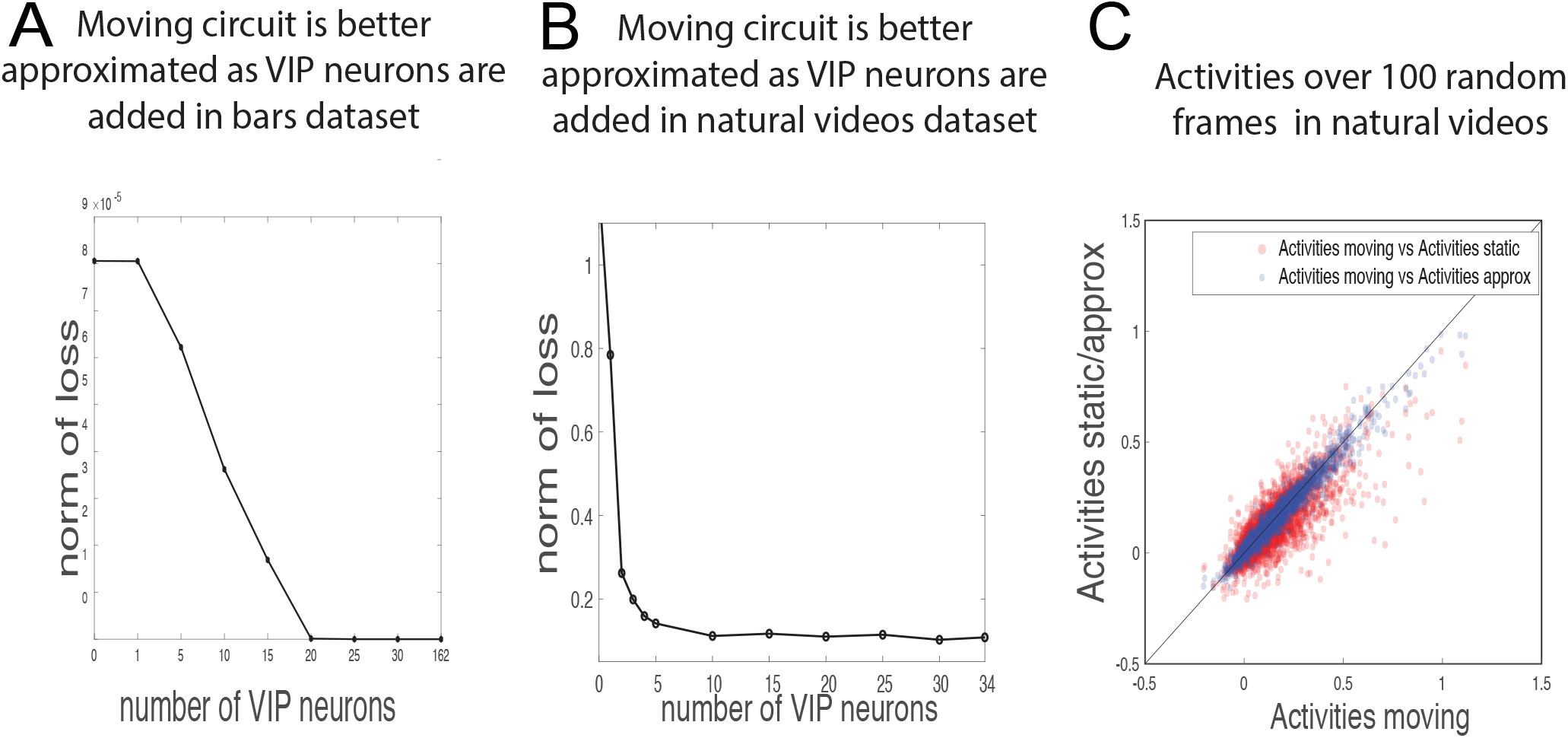
a. Adding VIP switching units to the circuit processing videos of bars approximates the activity to that of the optimal circuit for moving context for this simple dataset. However, no more than 20 VIPs are needed in practice, compared to the 162 PYR and SST cells. b. Adding VIP switching units to the circuit processing natural videos approximates the activity to that of the optimal circuit for moving context for the naturalistic dataset. However, no more than 5 VIPs per unit space are needed in practice, compared to the 34 PYR and SST cells per unit space. The parameters chosen for this optimization are Δ*t* = 2 and dim(**W**^*VIP→SST*^) = dim(**W**^*VIP→PYR*^) =34 × *Nf*_2_ × 3 × 3, dim(**W**^*PYR→VIP*^) = *Nf*_2_ × 34 × 3 × 3, where *Nf*_2_ is the variable number of VIP units. c. A random subset of activities corresponding to different video frames, filters, spatial positions for the static, moving, and approximated moving circuit. Red dots for activities for moving circuit (**r**^moving^) vs activities for static circuit (**r**^static^); blue dots for activities for moving circuit vs activities for approximated switching circuit (**r**^approx^). Activities are computed using weights with 5 VIP units/unit space. Activities chosen for the approximated switching circuit are able to better estimate the activities in the moving circuit in comparison to the ability of the activities in the static circuit to estimate the activities in the moving circuit.

For images and videos of natural scenes, the *movement approximation error* in (18) was minimized when the number of VIP units is 34 per unit space, which matches the number of units in the PYR and SST population. However, the approximation error was already significantly minimized with only 5 VIP units per unit space, without any significant improvement after adding more units (Fig. 7b). Varying the dimensionality of spatial components of the tensors (Fig. S4) we were solving for (**W**^*VIP→SST*^, **W**^*VIP→PYR*^, **W**^*PYR→VIP*^) and the synaptic delay Δ*t* for sparse weights **W** that account for patch independence, we obtained the same qualitative results. Our results also hold for non-sparse weights, as shown in Fig. S5. Fixing the number of VIP units to 5 per unit space, we find that the approximated firing rate of (17) matches **r**^moving^ compared to the **r**^static^ firing rates of a circuit without VIP units (Fig. 7c). We conclude that for the specific parameters chosen in Fig. 7b, the ratio of PYR to switching VIP units is 34/5 = 6.9, so that the switching operation requires relatively few units, a fact we return to in the context of the underlying biology below.

All in all, we have shown that a switching circuit with relatively few numbers of switching VIP units and appropriate feed-back connections can be implemented to achieve visual processing during the static and moving contexts, and for both a simple synthetic dataset of bars, and a biologically relevant dataset of natural images and videos.

### 2.6 Context-dependent visual processing with extra-classical receptive fields leads to denoising

According to our theory (Methods, sec. 4.1), the moving circuit achieves optimality of visual processing for videos, the static circuit achieves optimality of processing for static images, and we have found appropriate connectivities to and from a population of switching units — VIP — that can approximate either circuit in a model of V1, the *switching circuit*. We have however not yet assessed the perfomance of these circuits on specific visual processing tasks. We pursue this here for the task of denoising. Specifically, we ask how well (a) extra-classical receptive field contributions from the static or moving circuits (Fig. 5a) can improve reconstructions of noisy videos and (b) whether the switching circuit can achieve the same level of performance as the separately optimized moving circuit when processing videos.

To reconstruct a visual scene during movement, our brain uses information from the present, but also time-delayed surround information, both of which can be inaccurate or incomplete. We use **W**^moving^ to weigh the past surround information, as these weights encapsulate the cross-correlational structure between features of the past and the present, thereby informing which features are more or less likely. We note that, during motion, using **W**^static^ to weigh surround information may still be better that using no surround at all: if movement in the videos is slow enough, or Δ*t* is small, features are smooth and **W**^static^ and **W**^moving^ are highly correlated.

To apply our models to the task of denoising, we apply Gaussian white noise or salt and pepper noise *ξ* to the original frames *X* of the videos (Fig. 8c), and compute firing rates in the circuits in responses to the noisy frames *X + ξ*. The firing rates are expressed as:

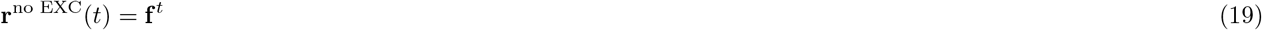

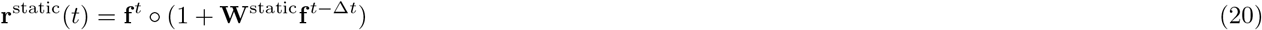

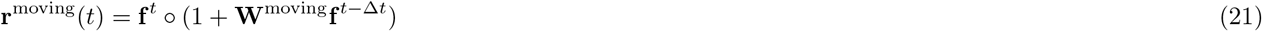

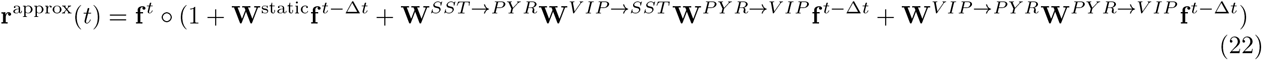

**Figure 8:**
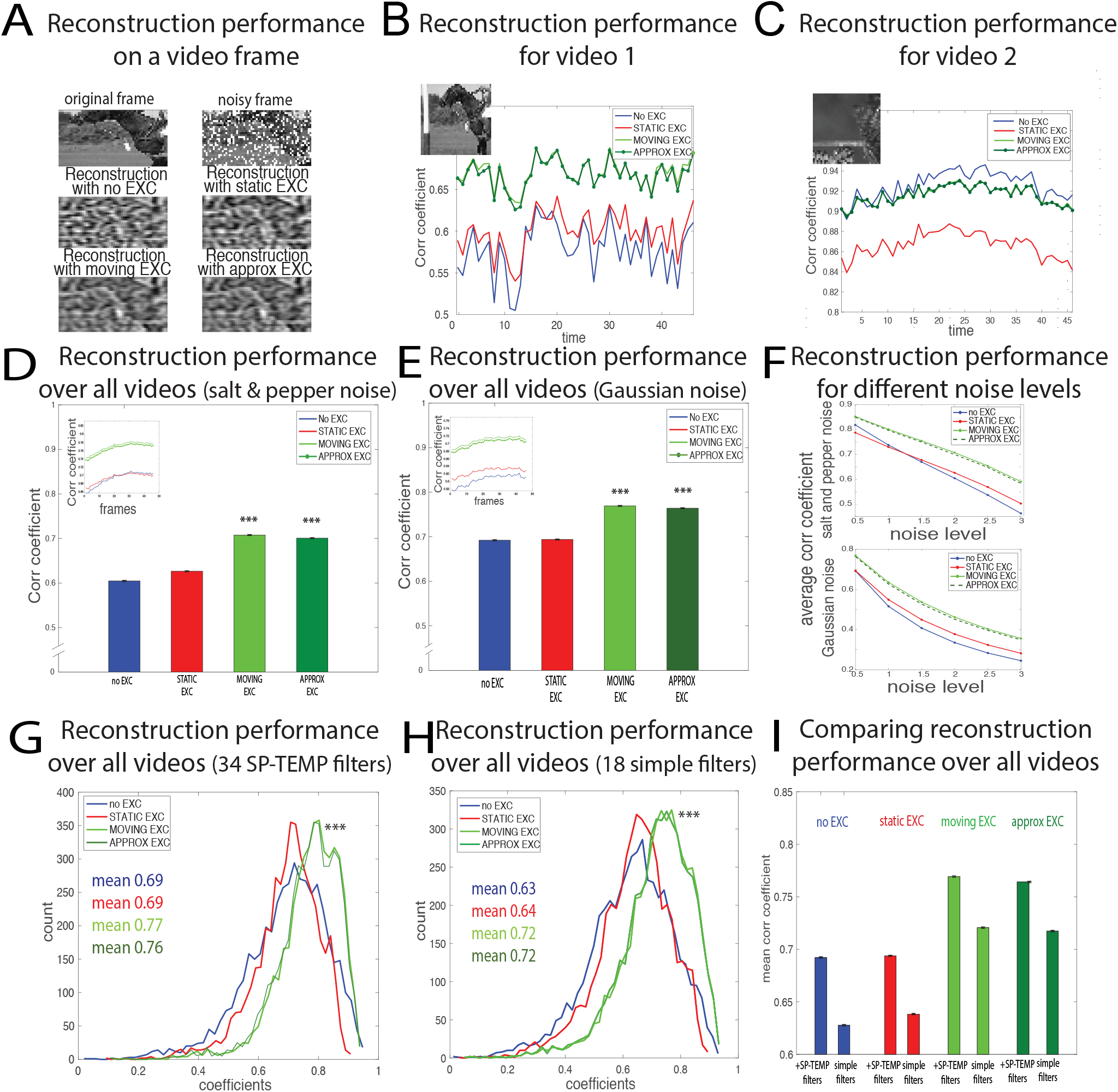
a. Example of a reconstructed frame for each condition/circuit architecture: no EXC, static EXC, moving EXC, approximated EXC. b. Average correlation coefficients between reconstructed noisy frames and reconstructed noiseless frames to assess denoising performance, for each frame in our video dataset. In this video, reconstruction benefits from surround contextual information. c. Same as a, but in this case the general inequality that holds on average *ρ*(**r**^no EXC^), *ρ*(**r**^static^) < *ρ*(**r**^moving^) ≈ *ρ*(**r**^approx^) breaks down and *r*(**r**^no EXC^) ≈ *r*(**r**^moving^). d. Average correlation coefficient over all frames and all videos after salt and pepper noise was added to the video frames. The probability is 0.2 each pixel is changed to white and 0.2 each pixel is changed to black. Δ*t* is set to 2 (frames). The average correlation coefficient is higher for moving and approximated EXC, than it is for static EXC, or in cases when no EXC is used (p-value < 0.05 using the Wilcoxon rank-sum test for all relevant comparisons). Inset: Correlation coefficients in time, averaged across videos. e. Same as d, for Gaussian white noise with 0.5 standard deviation. Δ*t* is fixed to 2 (frames). *p* < 0.05 for all relevant comparisons, Wilcoxon rank-sum test. f. Average correlation coefficient over frames and videos as noise level is varied. Top: Noise level as salt and pepper noise is varied; Down: Noise level as Gaussian white noise std is varied. g. Correlation coefficients over frames and videos for different conditions/circuit architectures when all 34 spatio-temporal filters are used. h. Correlation coefficients over frames and videos for different conditions/circuit architectures when only 18 filters are used. The filters used are ones without the temporal component. i. Comparison of average correlation coefficients across conditions/circuit architectures for the 34 spatio-temporal filters and the 18 “simple” spatial filters.

We denote “EXC” throughout the figures and text to represent the extra-classical receptive field contribution. Hence, **r**^no EXC^ is the firing rate due to only the feedforward pathway, with no lateral connections, and thus without any extra-classical, surround modulation. In the case of **r**^static^ (**r**^moving^), **W**^static^ (**W**^moving^) weights are the lateral connections applied that weigh the extra-classical receptive field information from the past surround. While **W**^static^ are non-optimal weights to compute the firing rate, **W**^moving^ are optimal for inferring features in noisy conditions as described below (cfr. Methods sec. 4.1). Finally, **r**^approx^ results from lateral connections from our switching circuit with connections to and from VIP.

For each image frame X we computed the corresponding firing rate **r** via equations (19) – (22), to obtain a tensor with entries for every filter and spatial position of X. We then deconvolved **r** for each filter **F**_*j*_ (Methods sec. 4.6) along its corresponding dimension to obtain the “reconstructed” frame *X′*:

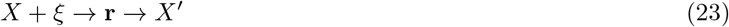

Although there are ways for a biological circuit to do more accurate reconstructions (e.g. via learning weights), we have chosen a simple reconstruction approach that does not require additional assumptions here (e.g. the circuit does not know the structure of the noise or the input), as described in Methods sec. 4.6.

We compare the quality of reconstructions from the four circuit models above. The baseline for these comparisons is the reconstruction of a noiseless image frame (*ξ* = 0), where the extra-classical contribution does not provide any additional information. (Note that this reconstruction *X′* is not the same as the original frame *X*, as all feature information not included in the filters is lost in initial convolution of the image frame to get **r**). We denote by *ρ*(·) a metric of the quality of the reconstruction. This takes the firing rate **r** as input, and generates the Pearson correlation coefficient between the reconstruction *X′* and the baseline reconstruction described above as output. The metric *ρ* for a video frame with noise *ξ* is

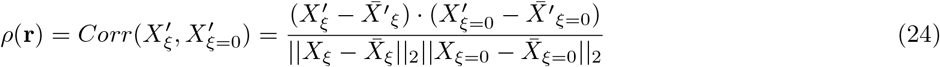

where · is the dot product, and 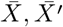 are the means of the image and reconstruction, respectively.

Thus equipped, we ask which circuit architecture gives rise to neural activity best suited for decoding visual scenes in noisy conditions. Fig. 8c shows reconstructions of a video frame using different such circuit architectures. We expect *ρ*(**r**^no EXC^), *ρ*(**r**^static^) < *ρ*(**r**^moving^), *ρ*(**r**^approx^) on average, as **W**^moving^ are the optimal lateral connections as defined above. However, the exact relationship between *ρ*(**r**^no EXC^), *ρ*(**r**^static^), *ρ*(**r**^moving^), *ρ*(**r**^approx^) depends on the exact correlational structure of the frames for each video. Some videos match our prediction that *ρ*(**r**^moving^) is maximized (Fig. 8a), while other videos do not (Fig. 8b). Specifically, there are videos where surround modulation is not effective, which appears to be due to the presence of independent features where the information in the extra-classical receptive field does not aid image reconstruction.

On average throughout the videos, **r**^moving^ and **r**^approx^ yield the best reconstructions (dark and light green bars), displaying the highest cross-correlation coefficients ρ between the noiseless reconstruction (the baseline) and the reconstructed frames (Fig. 8d). Figs. 8d and 8e show this holds true when we added to the original frames either salt and pepper noise, when we varied the proportion of pixels occluded, or Gaussian white noise, when we varied the standard deviation of the normal distribution of noise. The relation *ρ*(**r**^no EXC^), *ρ*(**r**^static^) < *ρ*(**r**^moving^) ≈ *ρ*(**r**^approx^) is robust to the amount of noise added to the frames (Fig. 8f), whether for salt and pepper noise or Gaussian noise. This holds true both when the complete set of 34 spatio-temporal filters is used (Fig. 8g), and when only the set of 18 filters with no temporal component is used (Fig. 8h). As expected, the addition of filters with a temporal component improves the reconstruction performance in all the four circuit architectures presented (Fig. 8i).

Thus, the switching circuit provides reconstruction performance comparable to that of a dedicated moving circuit. This is because the switching circuit reproduces firing rates that are close enough to **r**^moving^ to improve reconstruction fidelity. The correlation coefficients found between noiseless baseline reconstructions and reconstructions due to the moving and switching circuits, respectively, present almost perfect overlap (light and dark green curves in Fig. 8g, Fig. 8h). In sum, we conclude that the extra-classical receptive field contribution in the moving circuit and approximated switching circuit generates neural activity that can be decoded to produce more accurate frame reconstructions.

### 2.7 Experimental evidence of VIP role in movement-related visual coding

#### Activity

Published experimental findings already provide strong evidence that the VIP inhibitory population acts to modulate the visual circuitry in a movement dependent manner [48, 20]. Very recent results show that VIP neurons respond synergistically to stimuli moving front to back during locomotion, a conjunction expected during locomotion in a natural environment for mice, with a preference for low but non-zero contrasts [42]. Such an activity matches the one required in our models.

Additionally, we perform a small set of new analyses of experimental data in the context of our model. These draw both on the literature and on the Allen Brain Observatory [1], which contains in vivo physiological activity in the mouse visual cortex, featuring representations of visually evoked Calcium responses from GCaMP6-expressing neurons in selected cortical layers, visual areas, and Cre lines. The dataset contains calcium activations across multiple experimental conditions, and here we focus on periods of spontaneous activity, natural images, and drifting gratings.

Our model of the switching circuit shows that the relative number of VIP neurons required to switch between moving and static contexts is relatively low when compared the number of PYR or SST neurons (Figs. 7a to 7b). This number qualitatively matches the relative abundance of neurons in the three populations. Excitatory neurons PYR are more abundant than inhibitory ones (roughly 80% to 20%), and VIP are a minority of inhibitory cells. Moreover, the existing VIP cells recorded in the Allen Observatory do not appear to exploit substantially more degrees of freedom (as measured by their relative dimensionality) than other cell populations (Fig. S8a), consistent with a small number of effective VIP “units.”

We now highlight two aspects of VIP neural activity which are directly related to our model and which justify the choice of VIP as switching units whose activities are modulated by the locomotion state of the animal. First, VIP activity dimensionality is significantly modulated across the moving and static conditions during periods of spontaneous activity, as shown in Fig. 9a and Fig. 9b. To extract such dimensionality modulation, we considered periods of spontaneous activity in the recordings and divided the statistical distribution of the animal’s speed, for each experimental session, into 4 quartiles. We then computed the average *dimensionality*, or Participation Ratio (PR, cf. Methods sec. 4.7) for each recording in each quartile, which we define here as the (lower) dimension of a subspace where the data of activations can be represented while retaining some meaningful properties of the original data. We define the “dimensionality modulation” to be the ratio between the average speed distribution within the highest quartile (movement condition) and the average within the first quartile (static condition). Such ratio is displayed in Fig. 9b. The dimensionality of the VIP population is significantly modulated by movement, while in other populations the same quantity was not significantly different across moving and static conditions (Fig. 9a). The histogram of such statistics is shown in Fig. 9b.

**Figure 9:**
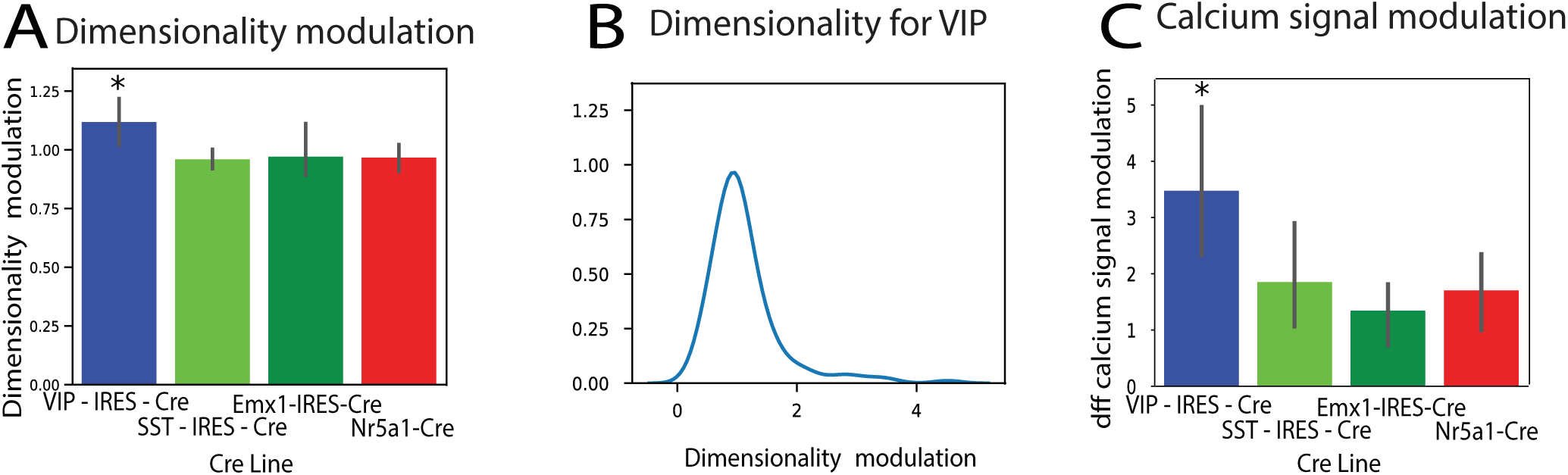
Data analysis of VIP population activity in calcium imaging data. a) Dimensionality ratio (Participation Ratio measure) during periods of spontaneous activity between movement and static conditions across CRE lines. b) Histrogram of the modulation of dimensionality (statistics relative to the blue bar in panel (a)). c) Activity (dff signal) ratio during periods of natural images viewing between movement and static conditions across CRE lines.

Second, we analyzed evoked activity during the animals’ viewing of natural scenes. We performed a Calcium signal modulation analysis and found that, for this stimulus set, the activity was strongly modulated for the VIP population and less so for other neural populations (Fig. 9c) across moving and static conditions assessed via the quartile method just described. This further confirms the stronger VIP modulation across the moving-static conditions. In the supplementary we discuss further pieces of experimental evidence, cf. Fig. S8.

#### Connectivity

While not as strong as the evidence regarding activity, we find connectivity data to be consistent with our model. Connection weights in the model can be interpreted as corresponding to a combination of connection probabilities and connection strengths in the data, as these have been shown to correlate well [12]. Connection probability as a function of the difference in orientation tuning (figs. S2c to S2d) qualitatively matches the same graph reported experimentally [31]. This like-to-like connectivity, with neurons responding to similar features (orientations) more strongly connected, holds true for both static (shown in [26] and figs. S2c to S2d) and moving weights (shown in Fig. S3). A second feature concerns the amplitude of static and moving weights which decreases with distance from the classical receptive field, with lower weights on average between neurons whose classical receptive fields are far away. Fig. S2 shows the dependence of the maximum, minimum, and average positive and negative synaptic weights, on distance between neuronal receptive fields. Assuming an exponential spatial decay of weights with distance and using the first two points in the plot displaying decreasing distance dependence in the mean positive static weights curve (Fig. S2a), we computed the spatial constants *D*_static/moving_ = 0.8× the classical receptive field size. This is in accordance with past findings [2, 26], suggesting that the near surround extends over a range which is similar in size to the classical receptive field.

Experimental data on connectivity in the visual cortex has shown that in layer 4 of V1, the average connection probability from VIP to SST is double the connection probability from VIP to PYR (0.625 compared to 0.351), while in layer 5, VIP to SST is 5 times more probable (0.625 compared to 0.125) [48]. VIP to SST connections are also stronger than VIP to PYR throughout all the layers (0.32 compared to 0.28) [48]. When we examine the weights W we have inferred in our model, we find that there are a few, equally correct solutions for the optimization problem (18) due to the multiple local minima of the movement approximation error. One of the possible solutions we found matched experimental data showing that in various layers of V1, the VIP to SST connection is strong compared to other connections, specifically the VIP to PYR connection (Fig. S7a). Interestingly, this property arose only when including weights from SST to VIP in the circuit, consistent with experiments (Pfeffer et al. [48] found the connection probability/strength to be quite strong between SST to VIP: 0.77 for connection probability, 0.5 for connection strength [48]). We conclude that our model can work well with strong weights from VIP to SST, making use of the observed disinhibitory motif (Fig. S7).

Altogether these comparisons provide further support for our modeling assumptions, and for the role of VIP neurons in visual coding across static and moving conditions. Further analysis of future datasets, as examined in the Discussion section, will guide next steps of circuit modeling.

## 3 Discussion

We have introduced a computational model for V1 circuitry that uses multiple cell types to integrate contextual information into local visual processing, during two different — static and moving — contexts. We have identified a need for recurrence, leading to the architecture of a *switching circuit* with bidirectional, learned connections to a switching population (here, the VIP cell class). Beyond V1 and biological circuit modeling, this circuit may be useful in searching for artificial neural network (ANN) architectures that can operate in different contexts and switch effectively between them.

Our model connects to a body of recent empirical studies elucidating V1 neural cell types and network logic. First, Niell and Stryker have established that as the speed of mice increases, the circuit increases spiking overall and changes the frequency content of local field potentials [44]. Potentially, distinct activity patterns during locomotion could be attributed to effects from eye movements, however Niell and Stryker [44] provide evidence against this hypothesis. These findings prompt us to model the network as a switching circuit that adapts its activity as the state of the animal changes from static to moving. Later studies have focused on the connection strengths for excitatory and inhibitory neurons: neurons display “like-to-like” connectivity [12, 31], whereby neurons with similar orientation tuning have a higher probability of connecting and display stronger connections on average. Pfeffer et al. describe the V1 circuit logic by using transgenic mouse lines expressing fluorescent proteins or Cre-recombinase, providing a consistent classification of cell-populations across experiments [48]. Three large non-overlapping classes of molecularly distinct interneurons that interact via a simple connectivity scheme were identified: PV, SST, and VIP inhibitory neurons. In particular, PV inhibit one another, SST avoid one another and inhibit all other types of interneurons, and VIP preferentially inhibit SST cells.

Another important development made by Fu et al. [20] has established that locomotion activates VIP neurons independent of visual stimulation and predominantly through nicotinic inputs from basal forebrain. This study was the first to propose the existence of a cortical circuit for the enhancement of visual response by locomotion, describing a modulation of sensory processing by behavioral state. These studies motivate us to choose VIP as switching units and to map the positive and negative weights of our model to connectivities between different neuronal populations. Finally, another study suggests that differentiated network response during locomotion can be advantageous for visual processing [15]: an increase in firing rates can enhance the mutual information between visual stimuli and single neuron responses over a fixed window of time, while noise correlations decrease across the population which further improves stimulus discriminability. The authors hypothesize that cortical state modulation due to locomotion likely increases visually pertinent information encoded in the V1 population during times when visual information changes rapidly, such as during movement.

There is a vast literature on models of efficient coding starting with Barlow 1961 [4], Attneave 1954 [3] (for a great description of this literature see Chalk, Marre, and Tkacik 2018 [11]). On one extreme, if the signal to noise ratio is high and additional constraints (e.g. sparsity) are introduced, such models emphasize redundancy reduction [46, 51, 24, 14, 5, 63, 16]. At the other extreme, if the signal to noise ratio is low, such models emphasize robust coding [29, 18]. We use a theoretical framework that emphasises robust coding and that we have selected because of its generality. It starts with an assumption on neuronal activation functionality (i.e. firing rates of neurons encode the probability of specific features being present in a given location of the image). This model describes local circuit interactions needed for integration of information from surrounding visual stimuli in noisy conditions for an arbitrary representation. The model matches multiple empirical findings, for example that statistical regularities of natural images give rise to “like-to-like” local circuit connectivities, as observed experimentally [12, 31]. However, in different contexts the model predicts different functional lateral interactions. Therefore, we looked at circuits which can implement multiple functional interactions in one circuit.

Our model also relates to other switching circuits reported in the experimental literature. For example, selective inhibition of a subset of neurons in central nucleus of the amygdala (CeA) led to decreased conditioned freezing behavior and increased cortical arousal as visualized by fMRI [23]. This therefore identifies a circuit that can shift fear reactions from passive to active. Another study has unraveled the cellular identity of the neural switch that governs the alternative activation of aggression and courtship in Drosophila fruit flies [32]. While these studies detail circuits responsible for switching behaviors, there are circuits switching between contexts: from *detection* of weak visual stimuli to *discrimination* after adaptation in mice [45]; from high response firing during active whisker movement, to low response when no tactile processing is initiated [65]; from odor attraction in food deprived larva switching to odor aversion in well-fed larva [61].

In contrast to this rich body of experimental studies, there are relatively few computational models proposed so far that explain switching of circuits [62]. We may compare our V1 circuit to the recurrent circuits utilizing FORCE learning, where a single unit or few units project their feedback onto a recurrent neural net and momentarily disrupt chaotic activity to enable training. VIP units in our model precisely resemble such output units providing feedback in the FORCE framework, but it is unclear how far this analogy goes and to what extent the framework in [56] is helpful in understanding V1 circuitry.

Another interesting example of circuit with flexible, context-dependent behavior has been proposed by Mante et al. in [38], where pre-frontal cortex (PFC) activity is modulated by the presence of a visual cue signaling which feature (color vs direction) the animals must integrate in a random-dots decision task. PFC functionality in this task has been modeled using a recurrent neural network (RNN) that takes the direction of motion, color of random dots, and visual cue as input, and outputs the appropriate, reward-generating, direction to saccade. This suggests the RNN enacts a potentially new mechanism for selection and integration of context-dependent inputs, with gating possible because the representations of the inputs and the upcoming choice are separable at the population level, even though they are deeply entangled at the single neuron level. The architecture of the model RNN proposed in this study is simpler than what we have laid out, while also attaining high flexibility. There are important differences between the framework outlined in this paper and our work: first, it is unclear what the number of weights in the network might be for the circuit in [38] to be multi-tasking. One of our main motivations has been to achieve a switching circuit with few added units and weights, so that the circuit has fewer weights to learn than two separate circuits processing the two contexts independently. It is unclear if this potential advantage holds in the case of Mante et al. Second, our circuit adapts to the statistics of both static and moving scenes and yields firing rates that are optimal for visual processing in either context. In the case of Mante et al., the circuit does not change momentary input processing when the context changes, it simply adapts its dynamics to integrate the appropriate feature and initiate the action that will be rewarded. Context takes on different meanings in these two instances: in our model, context is given by the statistical regularities of a certain environment, static or moving; in Mante et al. context refers to an input cue that changes the goals and reward dependencies of actions within the task. Importantly, we have focused on switching circuits that modulate their responses to different sensory contexts, as opposed to different input cues and behaviors. It is unclear whether identical or different mechanisms for switching apply in the case of sensory processing or action selection, when the animal changes scene statistics or behaviors, respectively.

Although our model is faithful to some aspects of the biology of V1 circuits, it has several limitations. First, it has been reported that during animal locomotion, firing rates of neurons more than double, at least in layers II/III of V1. Our firing rates are normalized to sum to roughly one across features and cannot reproduce a doubling occurring uniformly over features. Second, another study [15] reported that noise correlations are reduced during motion, but this does not occur in our model. Further, we model VIP as a switch which is off during the static condition and has an activation during locomotion dependent on input images, whereas data shows VIP activity is modulated at a finer scale and correlates strongly with speed [20]. In addition, VIP switching units in our model turn on based on perfect knowledge of whether the animal is static or moving, rather than based on more subtle time-varying visual or motor features. Furthermore, data from [31, 48, 28, 25, 33, 59, 10] on connection probabilities and strengths between neuron populations presents a richer, more complex picture than our simplified circuit. There is wide-ranging connectivity to and from PV, there are strong connections from PYR to SST in most layers, and the weights from SST to VIP are strong (in terms of both connection probability and strength across layers), details that our simplified model cannot describe. Enabling weights from SST to VIP showed that we can similarly infer weights to and from VIP so that we are able to approximate the circuit during the moving condition (figs. S6a to S6b). However, there are still many more potential connectivity structures between neuron populations our model does not describe.

From a computational perspective, our model makes several simplifications in describing context integration in circuits tuned to the statistical regularities of natural scenes. These include approximating a product with a sum in Equation (36) in Methods and ignoring higher order surround modulation going from Equation (30) to (32) in Methods. For simplicity, we have also limited the basis set of filters to one that extracts information about oriented edges in natural scenes. However, the computation of the extra-classical receptive fields need not be intrinsically limited to simple cells responding to Gabor-like filters, but can be extended to encompass neurons responding to more complex features in areas beyond V1. Switching circuits can occur more generally, including in somatosensory and auditory cortices, where some of the same neuronal populations interact using similar circuit logic [44, 6]. Populations of neurons in general switching circuits can respond to diverse stimuli (e.g. the VIP in auditory cortex are activated by punishment [49]).

Here, we showed how a biologically inspired switching mechanism can enable a network to efficiently process stimuli in two different conditions. Most artificial neural networks (ANNs) suffer from what has been termed “catastrophic forgetting”, by which previously acquired memories are overwritten once new tasks are learned. Conversely, humans and other animals are capable of “transfer learning”, the ability to use past information without overwriting previous knowledge. Proposed solutions to this problem, like elastic weight consolidation or intelligent synapses, are discussed in [30], [64], and [36]. When applied to a narrow condition of learning new contexts, our work adds a switching mechanism based on the connections among different cell types in V1. This may open new doors to artificial neural networks with analogous switching architectures.

## 4 Methods

### 4.1 A theory of optimal integration of static context in images

A theory of optimal context integration was first outlined in [26] and describes a probabilistic framework for inferring features at particular locations of an image given the features at surrounding locations. The probabilities of these feature occurring and co-occurring are then mapped to elements of a biological circuit (firing rates, weights).

#### Neuronal code

We assume the firing rate of neurons to be a function of the probability of a feature being present at a specific location of the image:

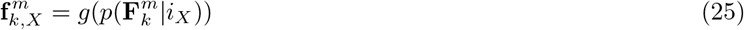

where 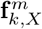 represents the firing rate due to the classical receptive field of a neuron coding for feature **F**_*k*_ at location *m* in response to image *i_X_*, and *g* is a monotonic function. For every image and every location we impose a normalization over features:

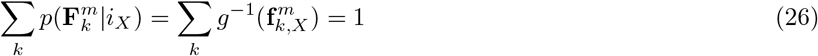

Thus, the sum over probabilities of features adds up to 1. Throughout the paper, we assume *g*(*y*) = *y*, although the model may be applied with other monotonic functions as well.

#### Probabilistic framework

We subdivide the image *X* into *N* patches that correspond to the classical receptive fields of neurons. Thus, we have:

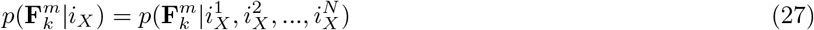

We will assume from this point forward that the firing rates are in response to an image *X*(*i_X_*), but omit the subscript *X* to simplify the notation.

We first look at the simple case where there are only 2 patches: the classical receptive field (patch *i^m^*) and the surround, which is part of the extra-classical receptive field (patch *i^n^*). We will take into account other surrounding patches later, when we perform an order expansion from 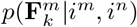 to 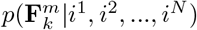. The aim in the simple case with two patches is to infer to what extent feature **F**_*k*_ at patch *i^m^*, denoted by 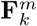, is present given information from both the classical receptive field and the surrounding extra-classical receptive field. Using Bayes rule and simple probabilistic relations, we sum over all possible features 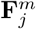 in patch *i^m^* to get:

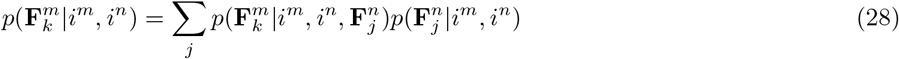

We can simplify the above relation by assuming the surround contribution from *i^n^* does not contain higher order surround information, instead it includes only data from the classical receptive field: 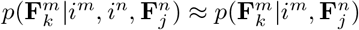. Our previous probabilistic statement (28) thus becomes

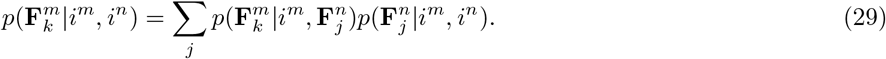

Using Bayes rule for the first term,

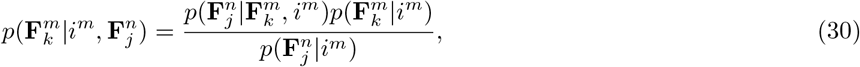

Equation (29) becomes

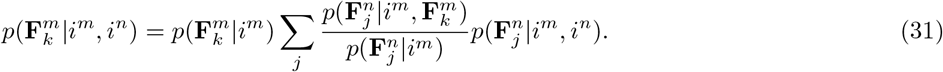

Assuming that we can ignore higher order contributions due to surround modulation, i.e. the surround modulation of the surround, we can make the following simplifications: 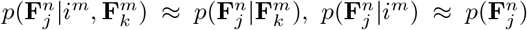, and 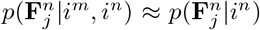. This way, patch *i^n^* is in the surround of patch *i^m^* and modulates the firing rate due to *i^m^*, but we are not concerned about the further effect *i^m^* has on *i^n^*. Then equation (30) thus becomes

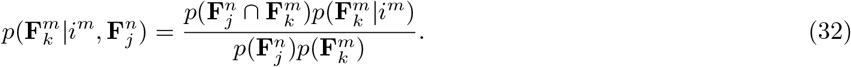

The original equation (28) becomes:

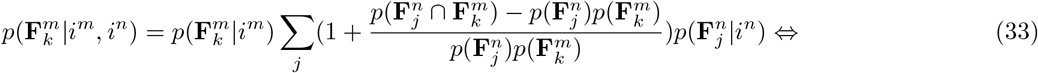

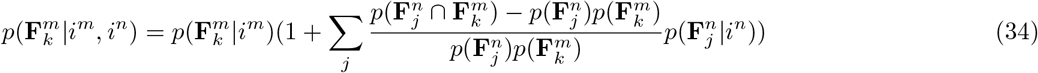

The last equivalence holds because we have assumed in (26) that all probabilities sum to 1.

We can now go from two patches to *N* patches that cover the entire image: *i*^1^, *i*^2^,…, *i^N^*. We further assume that each patch provides independent information to a neuron coding for 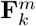 so that we obtain:

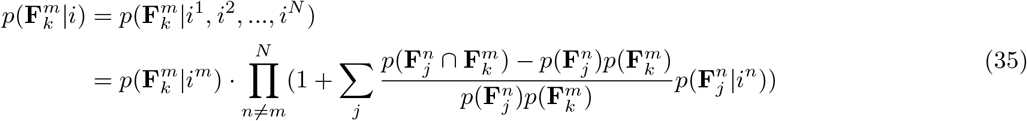

If the contribution from each patch is very small, we can ignore the higher order terms in (38) and apply the approximation 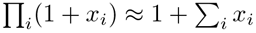 for *x*_*i*_ ≪ 1:

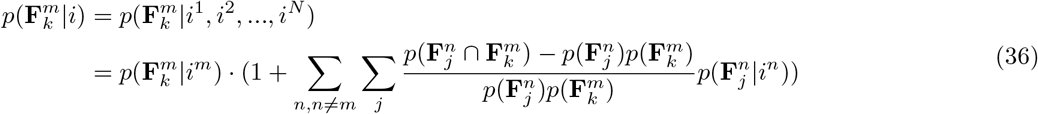

#### Mapping from the probabilistic framework to a neural network

Using a simple neural code with *g*(*x*) = *x*, so that the firing rate represents the probability of feature presence, we obtain a simple mapping to a network of neurons. We denote

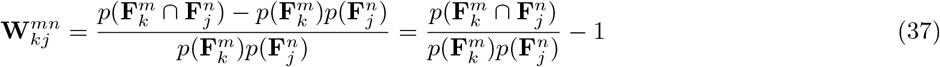

and map 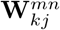 to the synaptic weight between neurons responding preferentially to features 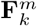 and 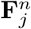, respectively. Then equation (36) becomes,

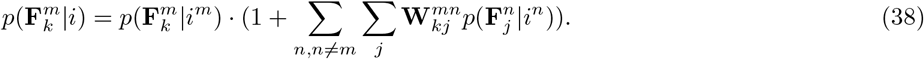

We can also map firing rates to probabilities: 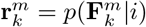 and 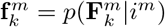, where 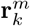 is the firing of the neuron with receptive field at patch *m* and most responsive to feature **F**_*k*_, and 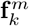 is the firing rate of the same neuron due to just the classical receptive field *i^m^*. As we recognize below, inferring these firing rates from our image and video datasets requires rectification and normalization so that **f** and **r** can be interpreted as probabilities.

The formula for synaptic weight can be expressed based on average activities of cells, when *X* spans a comprehensive set of natural images:

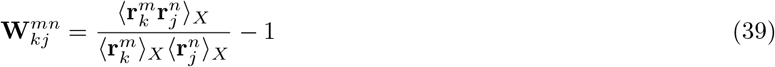

These weights can be achieved using Hebbian learning in an unsupervised manner. To avoid writing implicit equations for the firing rates which are difficult to solve, and to make the computation tractable in practice without requiring learning, we use an approximation that requires only **f**, the firing rates due to the classical receptive fields:

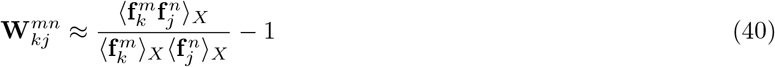

Finally, the probabilistic equations (36)–(38) outlined above can be re-written in terms of biologically-relevant quantities like firing rates and synaptic weights by applying the appropriate mappings:

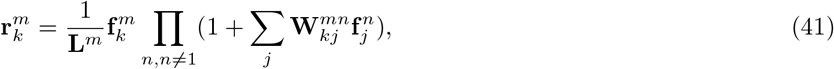

or, more simply,

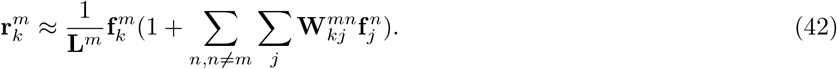

when lateral connections given by 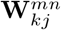 all sum up together to have a multiplicative effect. Here **L**^*m*^ is a normalization coefficient for patch *i^m^*, since we require

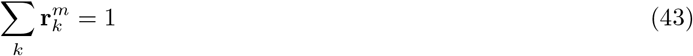

and thus denote

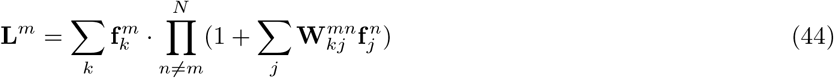

As outlined in [26], this can be implemented in a network in which a set of neurons responsible for normalization have a divisive effect on the neurons, are patch-specific (have a classical receptive field of similar size to the neurons), inhibit equally all the neurons in their image patch, are untuned to features in the visual space, and receive inputs equal to the average of the inputs of the neurons in the patch.

### 4.2 Computing the synaptic weights

To compute weights according to (40), we first compute 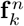, the firing rates due to the classical receptive field for every image *X* in a large dataset. Initially, we pre-process the image: we convert the image to grayscale, subtract the mean, and normalize the image to have a maximum value of 1. Similarly, we pre-process the filters so the mean of each is 0. **f**_*k*_ is the result of convolving *X* with feature *k*, rectifying and then normalizing so that at each location *n* the sum over features *k* of firing rates 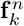 is equal to 1. Rectification ensures that firing rates are non-negative, while normalization further ensures we can interpret **f** as probabilities. We average these firing rates over all images *X* in the dataset to obtain 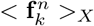 for each feature *k*. The feature co-occurrence probability given by 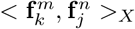 in the numerator for the synaptic weight formula is then computed by further pairwise convolution of firing rates due to the classical receptive field for each possible pair of filters in the basis set and each image in the dataset, and then averaged over all images.

For a dataset of videos, formula (40) becomes

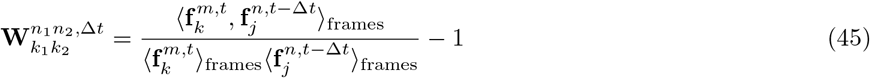

The feature co-occurrence probability given by 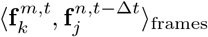 is computed by convolution of firing rates due to the classical receptive field at different frames (*t* and *t* – Δ*t*) for each video and averaged over all videos and video frames. The assumption here is that extra-classical effects are delayed by a time Δ*t* that corresponds with the time between movie frames or, biologically, corresponds to the synaptic delay.

We first assume translational invariance so that only the relative position of two filters is relevant: 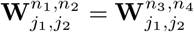 when 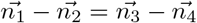. The assumption that weights act with translational invariance allows to rewrite the connectivities as simply a function of the distance, in image space, between the receptive field centers of the two neurons. Second, the mathematical validity of our probabilistic framework relies on the assumption that patches in the visual space, representing receptive fields of neurons, contain independent information. To reconcile this assumption with our empirically derived weights, we only consider connections between neurons whose receptive fields are sufficiently far apart, regardless of their corresponding feature identity. This leads to the usage of sparse weights for moving and static contexts (Fig. 4e), where the only non-zero weights we allow in W are spatially half of receptive field apart. More precisely, for every feature *k*, synaptic weights from target filters were sampled in steps of 0.5× the receptive field size at 3 distances in each direction around (0, 0), so that we have synaptic weights on a (7 × 7) grid (3 connections to the left/up + 3 connections to the right/down + self-connection = 7). Instead of using these sparse weights after sampling, we could have also re-scaled the original, non-sparse weights by a scalar *α* so that ║**W**^statlc/moving (sparse)^ – *α***W**^statlc/moving^║ ≈ 0. Searching over possible values of α, we find *α* ≈ 1/50. We choose however to work with sparse weights, or test our results on the original, non-sparse weights without worrying about the re-scaling by *α*. Although results presented in this study are largely for sparse weights, we have checked that the main results also hold when using full connectivity, at least for small Δ*t* ∈ {1, 2} (Fig. S5a). Further, assuming that the contribution due to context integration decays as the filters are spatially further and further apart, we can limit the weights in space to three times the size of the classical receptive field. Sample synaptic weights obtained using this procedure are shown in Fig. 4e (and Figures Figs. 4d and 4f without the sampling of weights).

### 4.3 Constructing the feature space for natural images and videos

We chose a basis of spatial filters that was constructed as outlined in [26]. This is done by averaging approximations of spatial receptive field sizes from 212 recorded neurons in V1 [19]. This set of filters is our first feature space and consists of four classes of spatial RFs observed experimentally: ON (1 feature), OFF (1 feature), and two versions of ON/OFF neurons (8 features each, for a total of 16), with the first version having a stronger ON subfield, and the second a stronger OFF subfield. Each subfield was modeled as a 2D Gaussian with a standard deviation of *σ* = 0.5× average subfield size, which was measured to be 4.8 degrees for the OFF subfield, and 4.2 degrees for the ON subfield. The relative orientation between two subfields for each ON/OFF class was varied uniformly in steps of 45 degrees, from 0 to 315 degrees. Also for the ON/OFF class, the relative distance between the centers of the ON and OFF subfields was chosen to be 5 degrees, which equates to roughly 2*σ*. According to the data, the amplitude of the weaker subfield is chosen to be half that of the stronger subfield, whose highest amplitude was chosen to be unity. These two subfields are then combined additively to form a receptive field whose size is 7 degrees (the distance between the two subfields plus *σ*). The set of 18 features is shown in Fig. 3d.

We then added 16 more filters with a temporal component, for a total of 34 filters. These filters have 2 frames with the first frame being one of the ON/OFF filters. The second frame is the ON/OFF filter in the previous frame shifted 3 pixels to the left, which matches the distance the sliding window moves every frame to generate the video. Such a spatio-temporal filter is shown in Fig. 3e.

### 4.4 Datasets of natural and synthetic images and videos

#### Natural images and videos

For the dataset of images, we used the Berkeley Segmentation Dataset (BSDS) training and test datasets [40]. The training dataset consists of 200 images of animals, human faces, landscapes, buildings etc. and is used compute the weights **W**^statlc^. This same training set is then employed to construct the dataset of 200 videos where a sliding window moves across the image for each frame of the video. In the simple case, the sliding window (167 × 167) moves 3 pixels per frame in the horizontal direction across the image (321 × 481 or 481 × 321), from left to right for 50 frames (Fig. 3b). The sliding window may also move in any random direction, resulting in different statistics of the video dataset and hence different **W**^moving^. This different dataset of videos is generated by choosing any pixel in the image and moving the sliding window toward it in smaller increments until that pixel is reached; a new pixel is then chosen from the image until there are a maximum limit of frames in the video (50 frames). Results from this different dataset are shown in Figs. S1 and S2. We further get 100 images from the BSDS test set to generate the corresponding 100 videos and use in the optimization problem. These video frames are provided as input to the optimizer that minimizes the loss functions *E*_*switch*,1_ and *E*_*switch*,2_ to find **w**^*α*^, **w**^*β*^ for *E*_*switch*,1_ and **W**^*VIP*→*SST*^, **W**^*VIP*→PYR^, and **W**^*PYR→VIP*^ for *E*_*switch*,2_. For both optimization problems we set 50 frames aside from these 100 videos to compute the generalization error during the minimization procedure.

In order to generate the figures in Fig. 8, another set of 100 videos generated from BSDS testing dataset is altered by adding Gaussian and salt-and-pepper noise of different parameters to each frame. The resulting noisy video frames are used to establish the ability of the switching circuit to do visual processing of stimuli with better reconstruction capability than the circuit implementing the static extra-classical receptive field or without extra-classical receptive field (Section 2.7). Gaussian white noise has standard deviation *σ* = 0.5 for reconstructions in Fig. 8e, while salt-and-pepper noise turns pixels black or white with probability *p* = 0.2 each, for reconstructions in Fig. 8d, figs. 8g to 8i. Parameters *σ* and *p* are varied (*σ* ∈ [0.5, 3], *p* ∈ [0.05,0.3]) in Fig. 8f.

#### Synthetic datasets of images and videos of horizontal and vertical bars

This simple synthetic dataset consists of 18 images of horizontal and vertical bars (9 horizontal, 9 vertical). Images are 9 × 9, each image having a bar at a different location. Videos consist of bars moving in any direction 1 pixel at a time: left or right (for horizontal bars), and up or down (for vertical bars).

### 4.5 Deriving an equation for PYR firing rate consistent with V1 circuit architecture

Let **f** be the firing rate due to the classical receptive field, **r** the firing rate incorporating extra-classical receptive field information, and *W^X→Y^* the weights between neuronal populations X, *Y*. We can write *approximated* expressions for firing rates of PYR, SST, VIP neurons at time t:

a. When there is no feedback connection from PYR to VIP

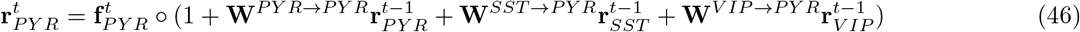

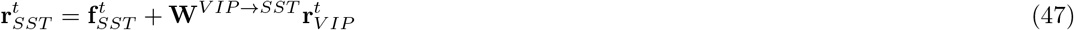

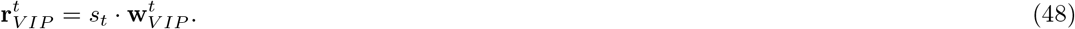
b. When there is feedback from PYR to VIP

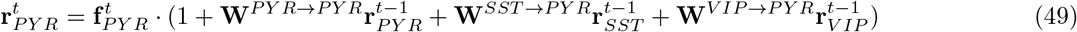

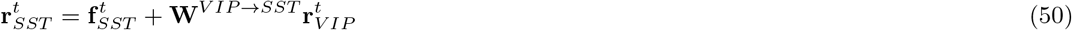

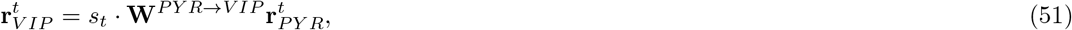

where *s_t_* is a binary variable that takes the value 1 during the moving condition and 0 during the static condition. For the analysis of the firing rate during movement we assume *s_t_* = 1. Equations (46) and (49), expressing the firing rate 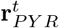 of the PYR population, assume the extra-classical receptive field contribution given by lateral connections has a multiplicative effect on the feedforward activities **f**_PY R_. This multiplicative gain is the result of mapping from the probabilistic framework of Equations (38) to (42) and their analogs for the moving circuit activities and weights. This results in the network doing optimal inference of visual features via PYR firing rates as expressed in (46) and (49), and as detailed in Section 2.1. The VIP firing rate **r**_VIP_ expression involves a binary gating term that switches based on state (static or moving), a simplification of what has been found empirically. The model could incorporate a term **f**^*VIP*^ into the expression (51) describing VIP firing rates driven independently from PYR such that 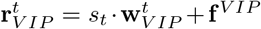, but this change would not alter our main results. Finally, only the inter-neuron connections with the longest synaptic delay are assumed to be non-instantaneous (connections to and from PYR), while other connections are presumed to occur at a much faster time-scale (connections between inhibitor neurons). Biologically, PYR are assumed to carry out computations by using dendritic trees, as outlined in [50], while SST and VIP are more spatially compact than PYR [22]. Hence, synaptic delays between PYR and other neuron populations are longer than between other populations.

Making the appropriate substitutions in (46) and in (49), we get the PYR firing rates:

for case **a)**,

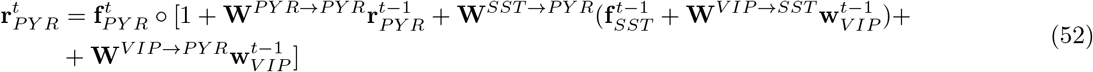

for case **b)**,

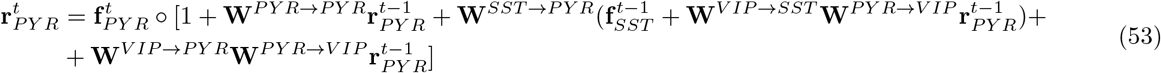

We can ignore further recurrence due to additional extra-classical receptive field contributions by making the approximation 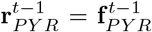. We are thus ignoring contextual surround modulation that is itself subject to surround influence — a “higher order” surround modulation — and instead consider only the classical receptive field response from surround neurons. These terms are small since this additional contribution is a linear combination of **f**_*i*_**f**_*j*_, **f**_*i*_**f**_*j*_**f**_*k*_, etc, where **f**_*i*_ are classical receptive field firing rates of neuron *i* and 0 ≤ **f**_*i*_ ≤ 1.

Additionally, we assume PYR and SST receive the same input so that 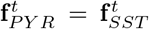. With these simplifications and dropping the subscript PYR for clarity, the equations for 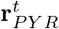 become:

for case **a)**,

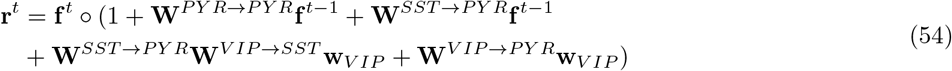

which leads to

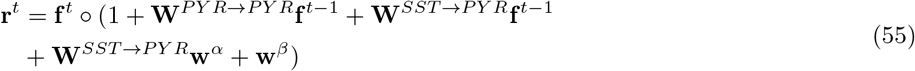

where **w**^*α*^ ≡ **W**^*VIP*→*SST*^ **w**_*VIP*_ and **w**^*β*^ ≡ **W**^*VIP*→*PYR*^ **w**_*VIP*_, while

for case **b)**,

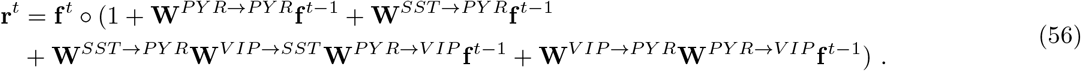

During the static condition, there is no contribution from the VIP and **f**^*t*^ = **f**^*t*–1^ so the firing rate becomes

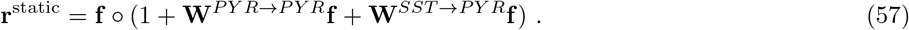

However, we know from our theoretical framework that the firing rate during the static context can be written as:

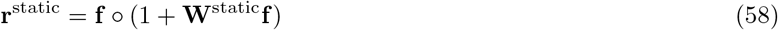

where **W**^statlc^ has been computed from the dataset(s) of images and is proportional to the average feature co-occurrence probability for pairs of spatial features. Therefore, we can consider a simple mapping that assigns **W**^*PYR*→*PYR*^ and **W**^*SST*→*PYR*^ to known weights: 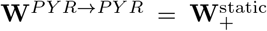 and 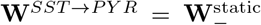, where 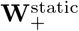 is the positive and 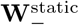 is the negative component of **W**^statlc^. The unknowns of equation (59) corresponding to the V1 circuit model with PYR to VIP connections, are thus only three sets of weights to and from VIP: **W**^*VIP*→*SST*^, **W**^*VIP*→*PYR*^, **W**^*PYR*→*VIP*^.

Finally, the equation for the firing rate of PYR neurons during the moving condition that we focus on throughout the paper (with PYR projecting to VIP) becomes:

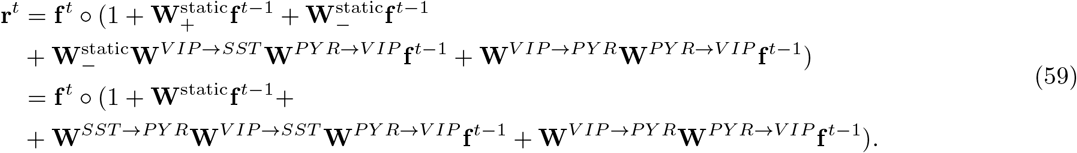

### 4.6 Reconstructions from noisy videos using firing rates and optimal synaptic weights of different circuit architectures

To gain insight into how optimal synaptic weights can facilitate decoding of information present in the neuronal activity, we reconstructed natural image frames from videos using 4 distinct circuits. The firing rates in these circuits are described by the following equations:

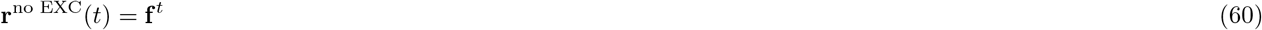

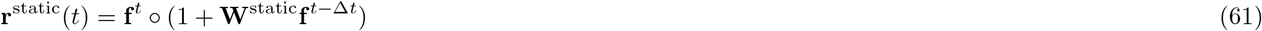

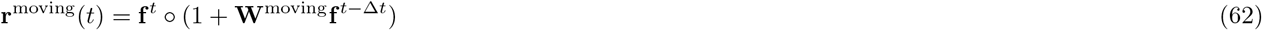

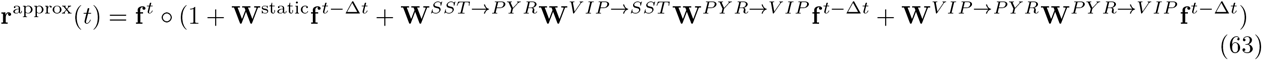

The first equation above describing **r**^no EXC^ relies solely on the feedforward information where no extra-classical receptive field contribution is included. The next two expressions re-state how the firing rates for the static and moving circuits require contributions from the extra-classical receptive fields through lateral connections **W**^static^, **W**^moving^, reflective of the statistical regularities of images/videos. Equation (63) describes the switching circuit we have implemented and characterized above and should approximate the firing rate in the moving circuit when VIP are active: **r**^moving^ ≈ **r**^approx^.

The reconstruction was performed as follows. For any noisy input image *X* + *ξ*, where *ξ* is some random variable representing a noisy process, we calculated the effective firing rate (activity) **r** of neuron/feature *k* at location *n* using the eqs. (60) to (63) above. To reconstruct image frames from firing rates, we convolved the firing rates computed with the inverses of the filters in our basis set. More specifically, the activity **r**_*k*_ corresponding to filter *k* was convolved with the inverse of *k*, which was obtained by flipping *k* about the horizontal and vertical axes. These convolutions for all filters were then averaged to obtain the final reconstruction.

We then performed the reconstruction for the same image frame *X* without any noise added. We assessed the de-noising capability of our circuits by computing the Pearson correlation coefficient *ρ* between the reconstruction of *X* + *ξ* and the reconstruction of *X*. The latter is a baseline for our comparisons, as there is no noise to remove from the image frame through extra-classical surround modulation. The Pearson correlation coefficient *ρ* is a function of the activity **r** of different circuit architectures and is discussed and compared across circuits in Section 2.7.

There are two further issues that merit further discussion. First, if the spectral content of the noise and image frame is known, a Wiener de-convolution can be applied which minimizes the mean square error between the estimated reconstruction and the original frame. Such a Wiener de-convolution would minimize the impact of de-convolved noise at frequencies with poor signal-to-noise ratio. However, we assume here that interpretation of signals is done without access to knowledge of this spectral content, but rather implementing a naive reconstruction as would be optimal in the noise-free limit. Second, given the presence of extra-classical surround contribution, the de-convolution operation may be more complex than the simple, filter by filter, convolution with the inverse filter **F**^*T*^. Specifically, the inverse may contain information about the cross-correlation of features. Again we work in the simplifying limit in which this is not the case. We do not exclude however the possibility that the biological circuit may apply a more complex reconstruction (e.g. via learning weights), an interesting avenue to explore in future work.

### 4.7 Measuring dimensionality with the participation ratio

We aim to characterize the dimensionality of the distribution of population vector responses representing neural activity. Across many trials, these population vectors populate a cloud of points. The dimensionality is a weighted measure of the number of axes explored by that cloud:

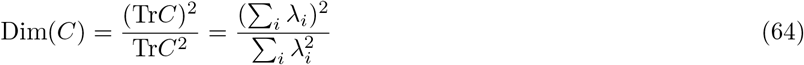

where *C* is the covariance matrix of the matrix of neural activations, and λ_i_ is the *i^th^* eigenvalue of the covariance matrix *C*. Dim(*C*) measures the dimensionality of neural activity of our network and is termed the *participation ratio*. The eigenvectors of the covariance matrix *C* are the axes of our cloud of points representing activity in neural space. If the neural activities are independent and all have equal variance, all the eigenvalues of the covariance matrix have the same value and Dim(*C*) = *N*. Alternatively, if the components are correlated so that the variance is evenly spread across *M* dimensions, only *M* eigenvalues would be nonzero and Dim(C) = *M*. For other correlation structures, this measure interpolates between these two regimes and, as a rule of thumb, the dimensionality can be thought as corresponding to the number of dimensions required to explain about 80% of the total population variance in many settings [41, 21, 34].

## 5 Supplemental figures

**Figure S1:**
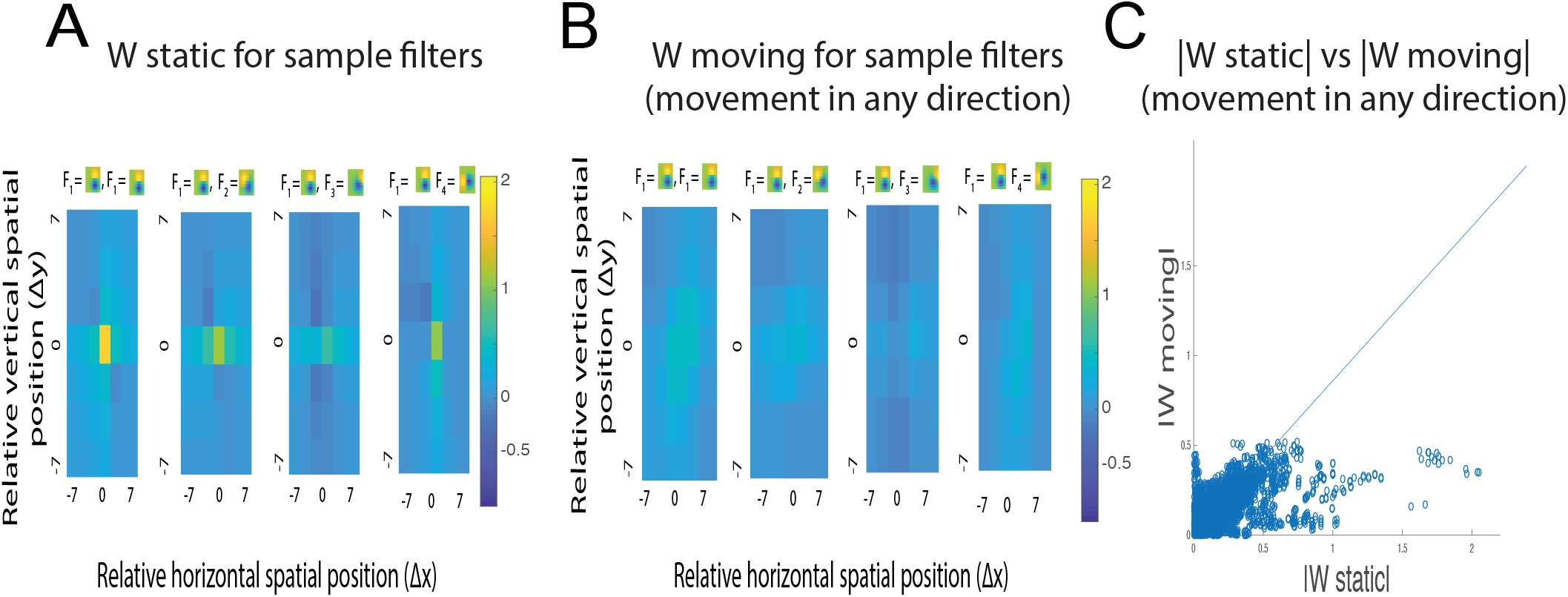
a. Slices of **W**^static^ corresponding to different pairs of filters (feature **F**_1_ paired with features **F**_1_ - **F**_4_). b. Slices of **W**^moving^ computed for dataset of videos where movement is in any direction. Slices shown correspond to different pairs of filters (feature **F**_1_ paired with features **F**_1_ - **F**_4_). c. Scatter plot of |**W**^static^| vs |**W**^moving^|. This reveals that on average, ║**W**^static^║ > ║**W**^moving^║ for this dataset of natural images and videos where movement can be in any direction.

**Figure S2:**
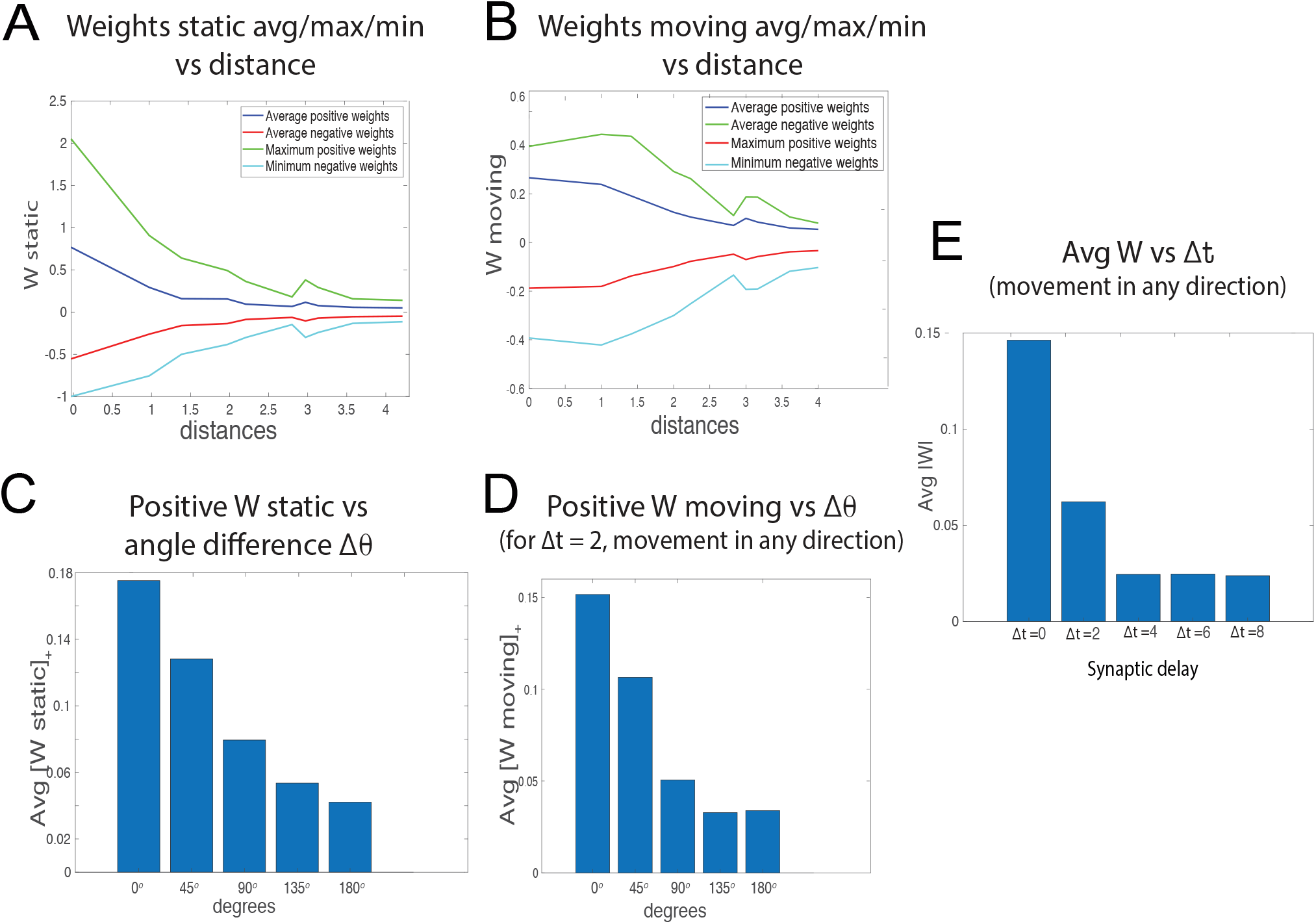
a. Dependence of the maximum, minimum, average positive and negative synaptic weights for the *static* context onto a target neuron *k* from all neurons on the distance measured in terms of receptive field size (1 unit = 1/2 RF size = 7 pixels). This distance dependence enables us to compute the spatial constant in terms of the classical receptive field size and compare it to data. b. Dependence of the maximum, minimum, average positive and negative synaptic weights for the *moving* context (Δ*t* = 2) onto a target neuron *k* from all neurons on the distance measured in terms of receptive field size (1 unit = 1/2 RF size = 7 pixels). The dataset of videos used to compute the weights here and in d, e is the one where the movement can be in any direction. c. Predicted average positive synaptic weight in the static context as a function of difference in orientation of features. This predicts that excitatory weights between neurons responsive to more similar features (similar in orientation) are stronger than those between neurons responsive to different features. The trend matches data in [12]. d. Predicted average positive synaptic weight in the moving context (Δ*t* = 2) as a function of difference in orientation of features. e. Average strength of moving synaptic weights as a function of Δ*t*, a parameter describing synaptic delay. The higher the synaptic delay, the closer to chance the co-occurrence probability is, and thus the lower the absolute values of the synaptic weights are.

**Figure S3:**
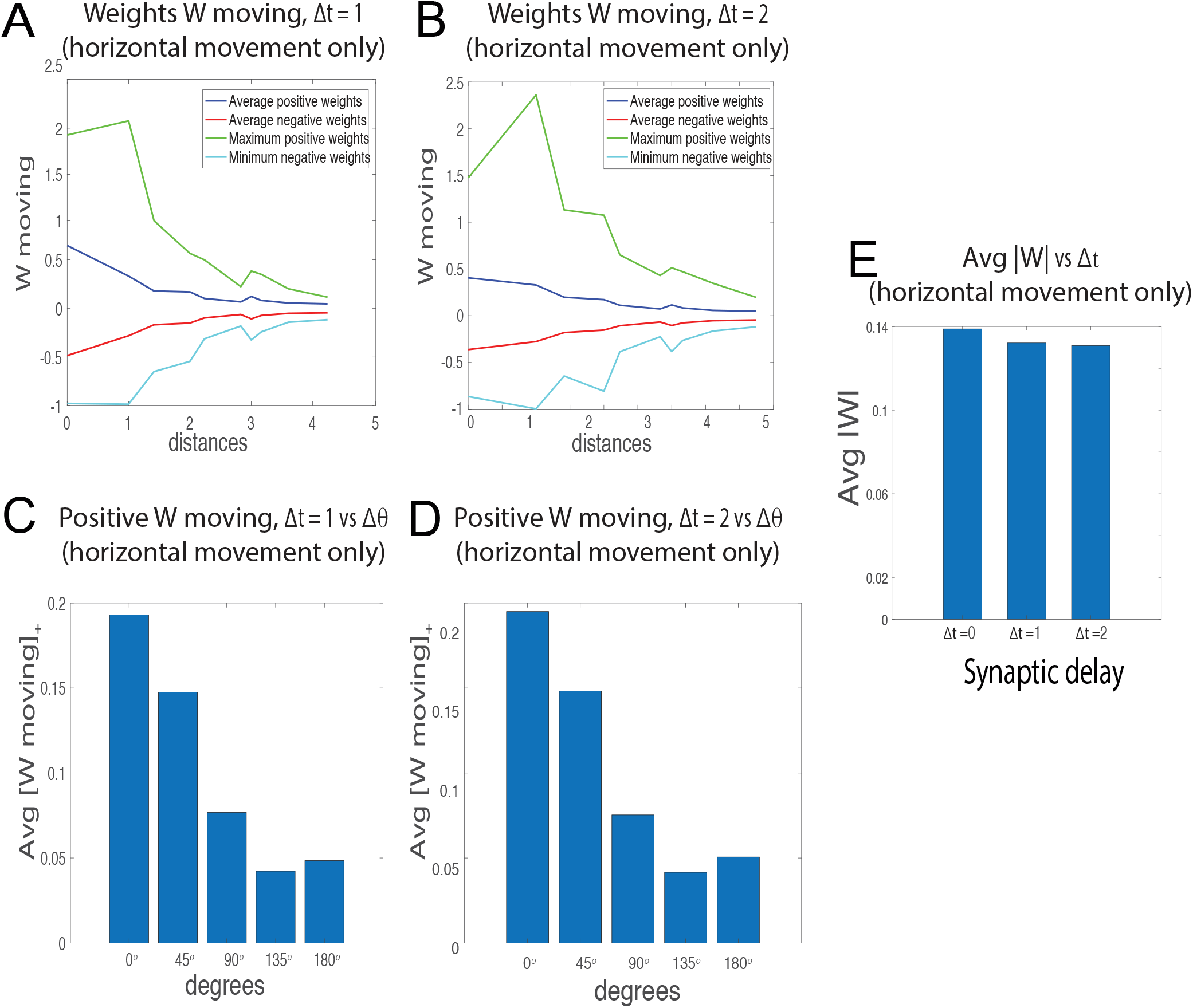
a. Dependence of the maximum, minimum, average positive and negative synaptic weights for the *moving* context with Δ*t* = 1 onto a target neuron *k* from all neurons on the distance measured in terms of receptive field size (1 unit = 1/2 RF size = 7 pixels). The dataset of videos used to compute the weights here and throughout this figure is the one where the movement can be only in the horizontal rightward direction. Because the movement is 3 pixels/frame and Δ*t* = 1 frame, the peak weight is between neurons responding preferentially to identical features and classical receptive fields centered 3 pixels apart (i.e. peak is at 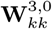, where Δ*x* = 3 ≈ 1/4 RF = 1/2 unit distance, not shown in the plot). b. Dependence of the maximum, minimum, average positive and negative synaptic weights for the *moving* context with Δ*t* = 2 onto a target neuron *k* from all neurons on the distance measured in terms of receptive field size (1 unit = 1/2 RF size = 7 pixels). Because the movement is 3 pixels/frame and Δ*t* = 2 frames, the peak weight is between neurons responding preferentially to identical features and classical receptive fields centered 6 pixels apart (i.e. peak is at 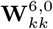, where Δ*x* = 6 ≈ 1/2 RF = 1 unit distance). c. Predicted average positive synaptic weight in the moving context (Δ*t* = 1) as a function of difference in orientation of features. d. Same as c, but with Δ*t* = 2. e. Average weight strength in terms of synaptic delay Δt, where Δ*t* = 0 corresponds to **W**^static^. Unlike the weights in Figure Fig. S2, which correspond to movement in any direction, the average weight strength does not decrease significantly with Δ*t*. Indeed, the peak of the tensor simply shifts at different spatial positions depending on how large the synaptic delay is, but otherwise the tensor remains (mostly) unchanged.

**Figure S4:**
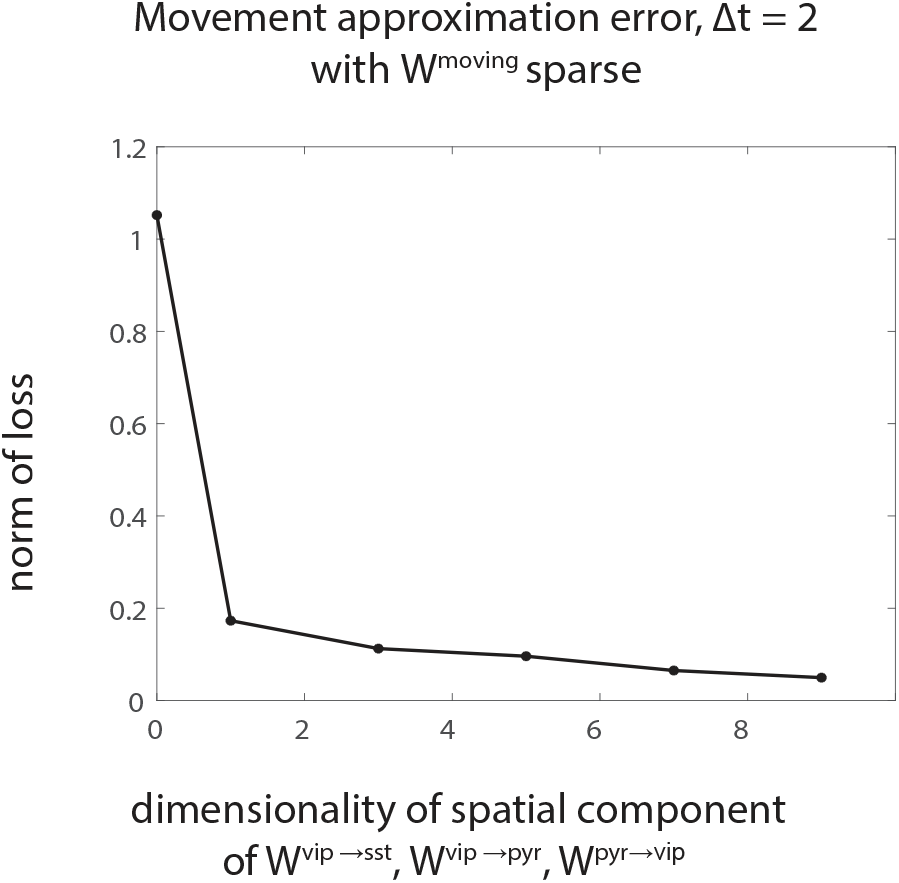
Varying the dimensionality of the tensors **W**^VIP→SST^, **W**^VIP→PYR^, **W**^PYR→VIP^ can lower the movement approximation error as defined in (18). These tensors have dimension *Nf*_1_ × *Nf*_2_ × *c* × *c*, where *Nf*_1_, *Nf*_2_ represent the number of VIP, SST, or PYR neurons, and c represents the dimensionality corresponding to the spatial component (shown on x-axis). We set Δ*t* = 2, *Nf*_1_ = 5, *Nf*_2_ = 34 for **W**^VIP→SST^, **W**^VIP→PYR^, *Nf*_1_ = 34, *Nf*_2_ = 5 for **W**^PYR→VIP^, and use sparse weights for the optimization procedure.

**Figure S5:**
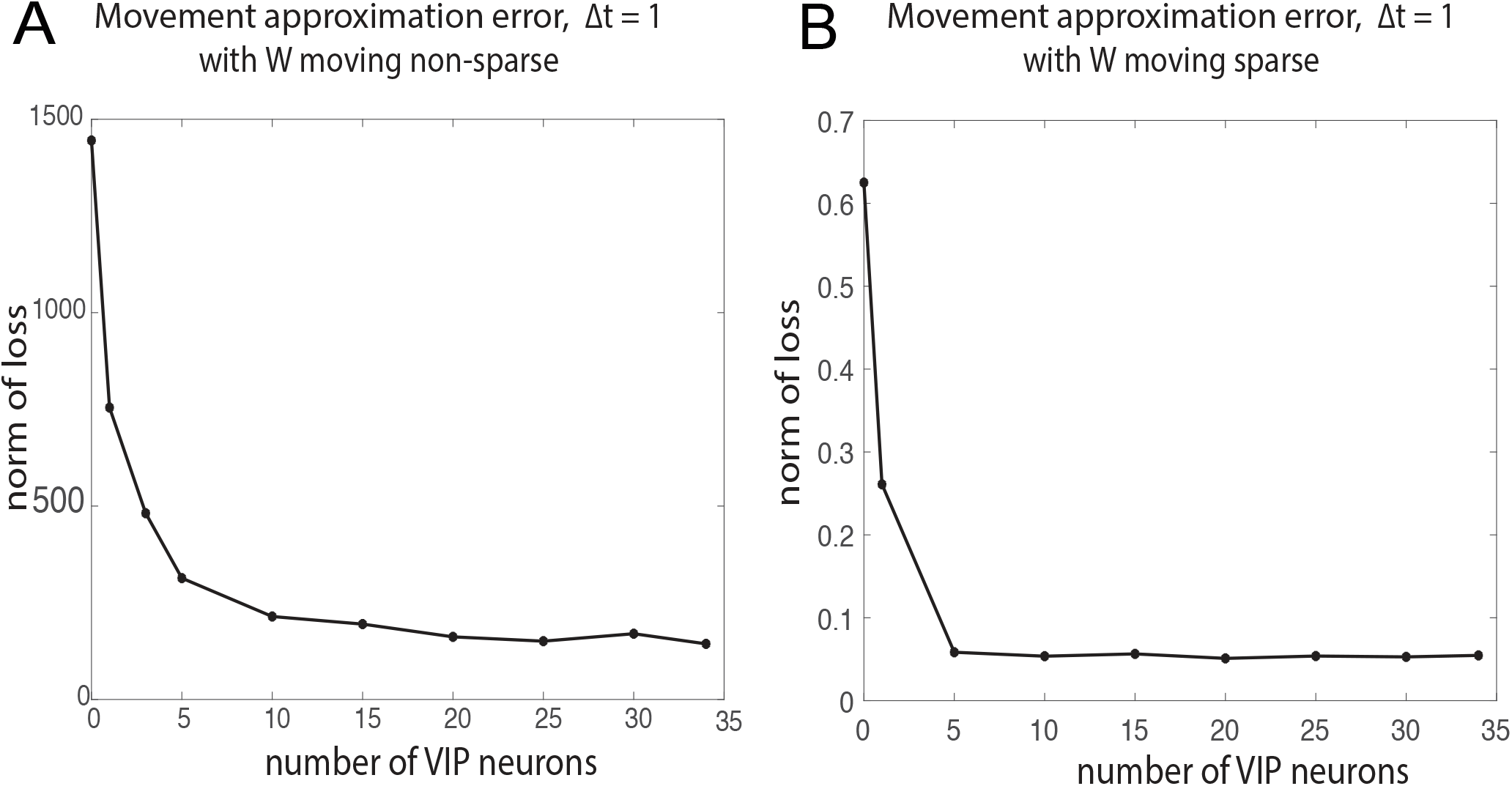
a. Movement approximation error (defined as in (18)) decreases with increasing number of VIP neurons for synaptic delay Δt = 1 and using the full **W**^moving^ (non-sparse). b. Movement approximation error decreases with increasing number of VIP neurons for synaptic delay Δ*t* = 1 and using the sparse sampled **W**^moving^.

**Figure S6:**
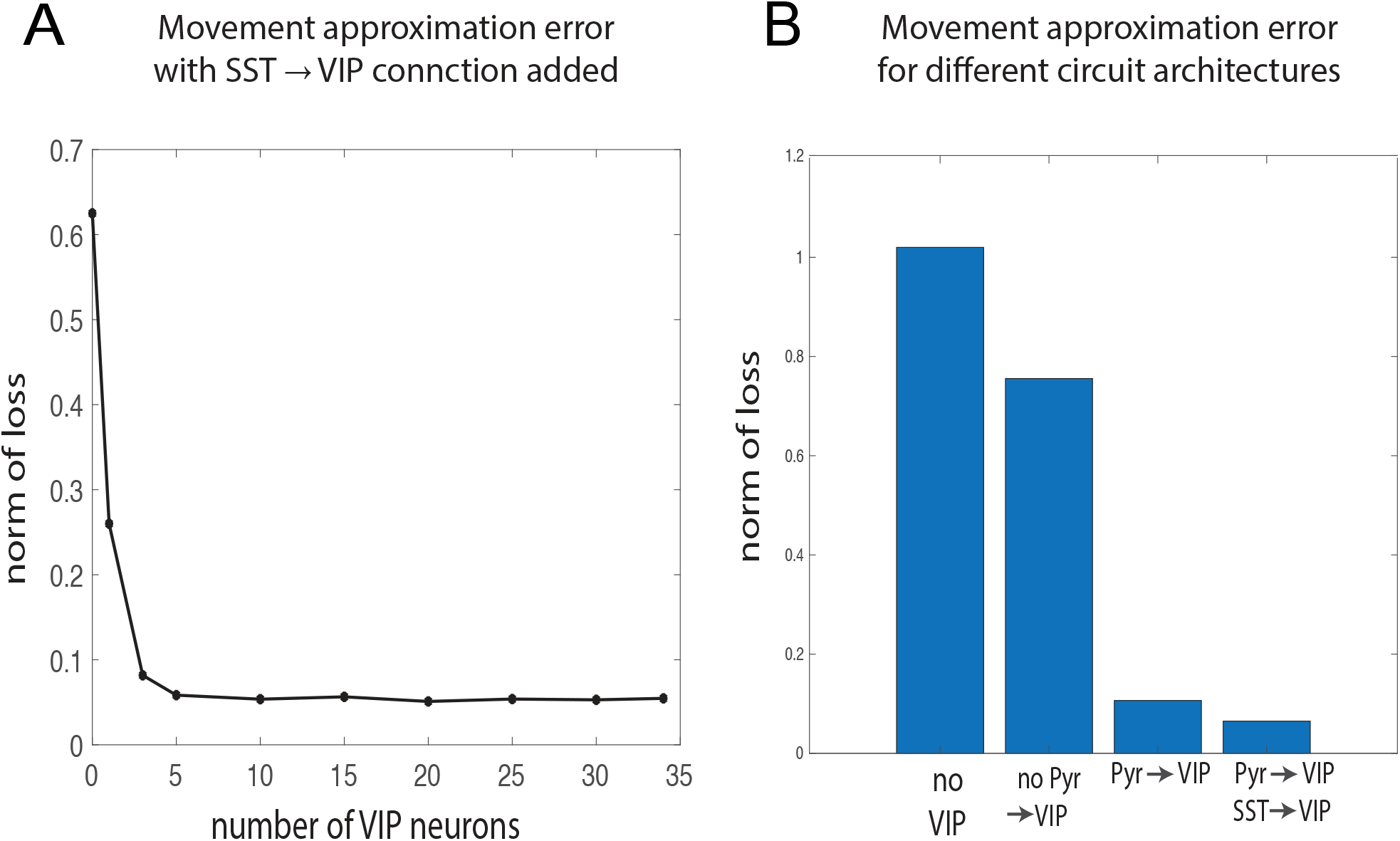
a. Movement approximation error (defined as in (18)) decreases with increasing number of VIP neurons, after an additional connection from SST to VIP is added. We set synaptic delay to Δ*t* = 1 and use the sparse sampled **W**^moving^. b. Movement approximation error for different circuits: a circuit with no VIP units (leftmost bar), a circuit with VIP and connections from VIP to PYR and SST (middle left bar), a circuit with an additional connection from PYR to VIP added (middle right bar), a circuit with an additional connection from SST to VIP added (rightmost bar).

**Figure S7:**
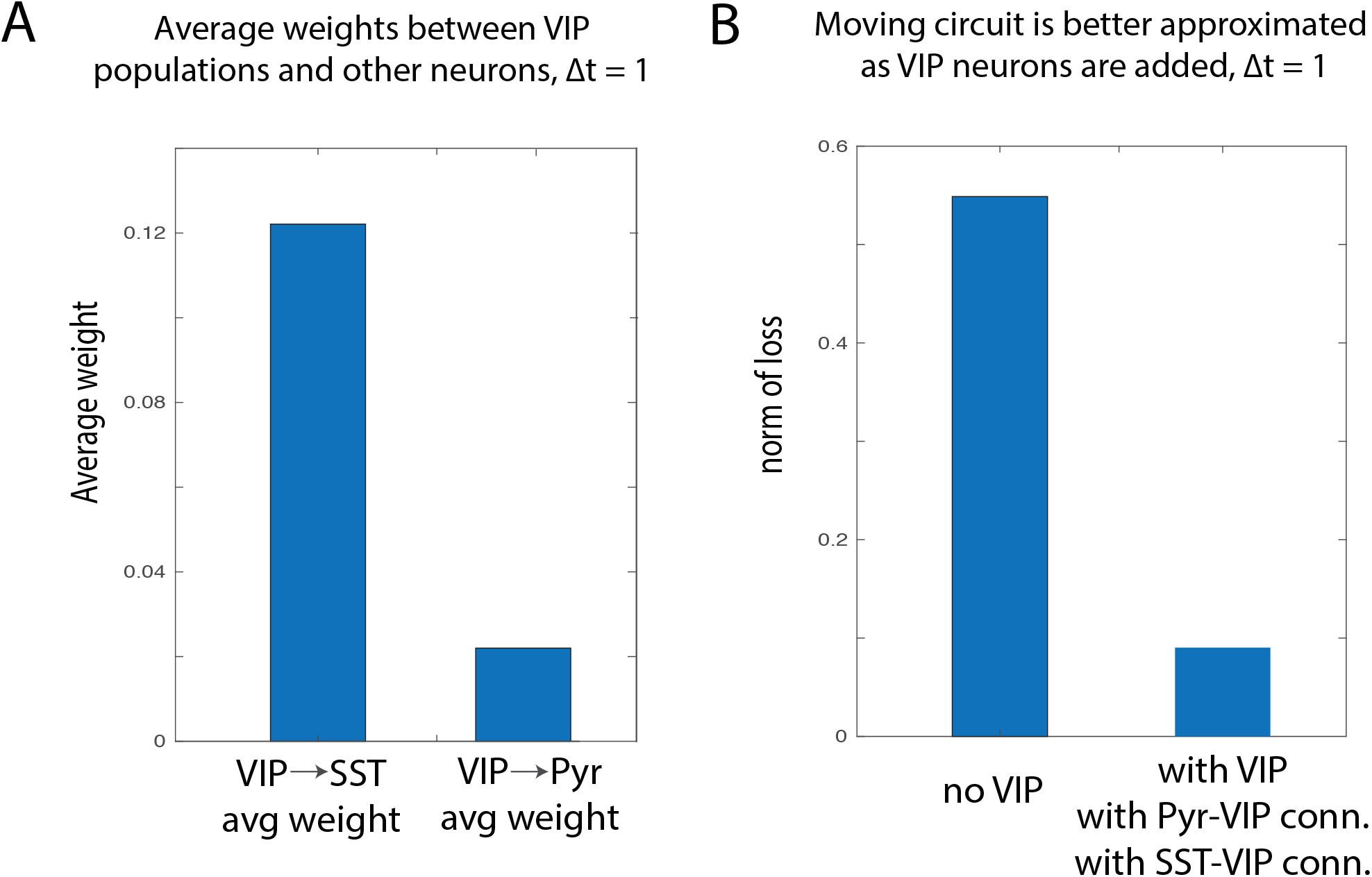
a. Comparison of **W**^*VIP→SST*^ average weights to **W**^*VIP→PYR*^ average weights (0.12 compared to 0.022). The ratio between these average weights is invariant to re-scaling due to patch independence that results in sparse weights **W**^*VIP→SST*^, **W**^*VIP→PYR*^. These weights have been computed by optimizing (18) for **W**^moving^ with Δ*t* = 1 (although a similar result holds for Δ*t* = 2) b. Verifying that using the solutions **W**^*VIP→SST*^, **W**^*VIP*→*PYR*^ to the optimization problem (18) yields a small movement approximation error (right bar) compared to the same error *E*_*switch*, 2_ when no VIP units are considered (left bar). The movement approximation error when VIP units are added (right bar) is for the circuit that includes SST to VIP and PYR to VIP connections.

**Figure S8:**
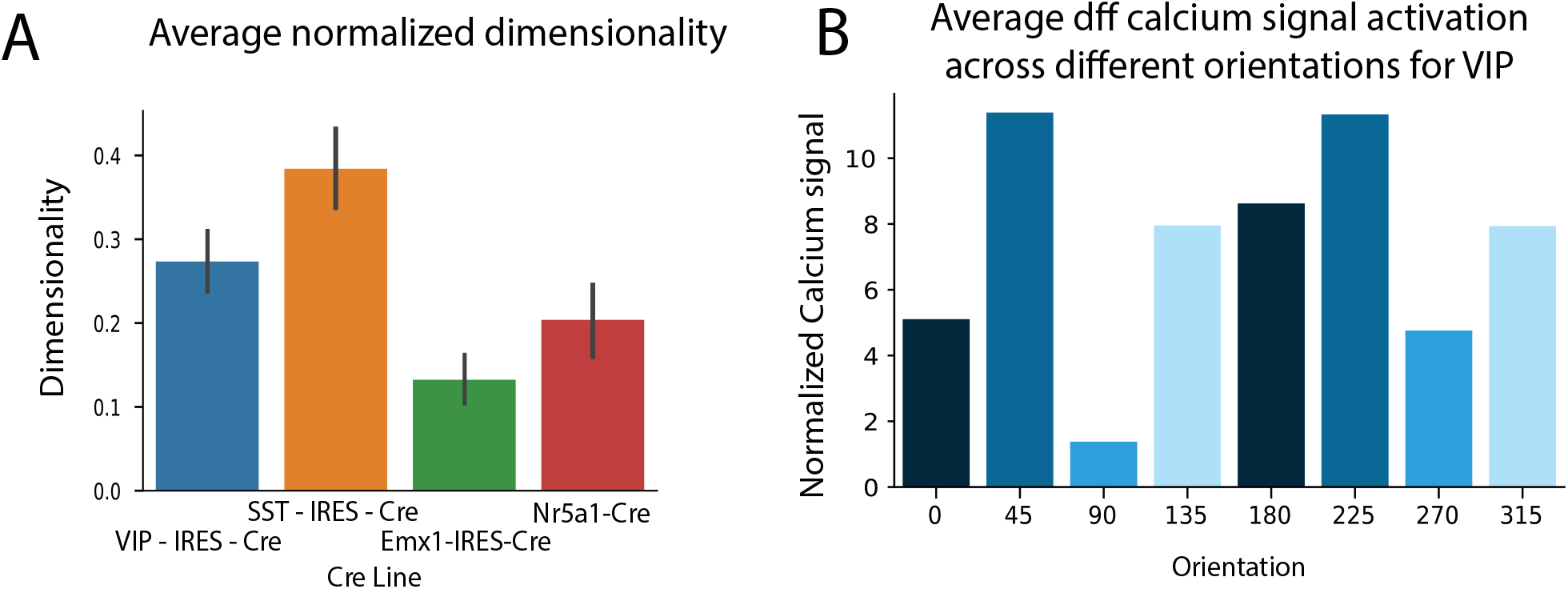
a. Average dimensionality across sessions normalized to the number of neurons in each session for multiple neural populations. Dimensionality is assessed by means of the measure Participation Ratio during epochs of spontaneous activity for the dff signal of calcium. While the average dimensionality of the activity of the PYR population is lower, this is partially due to the number of PYR units recorded being higher. b. Average dff calcium signal activation across different orientations for the VIP population during drifting gratings stimuli. Despite the trend appearing across orientations this is not significant as the Standard Error (not shown) is high due to the high variability across recordings.

## Footnotes

1 In general, if *N* is the number of neurons per location, *L* is the number of locations, and *C* is the number of connections per neuron, then the total number of connections in a circuit is *NLC*. Two identical circuits have *2NLC* connectivities, while a switching circuit has *NLC* + *LM*(*c_in_* + *c_out_*), where *M* is the number of switching (VIP) units, and *c_in_, c_out_* are the number of connections to and from the switching units, respectively. When *M* ≪ *N*, then *c_in_, c_out_ < C* and thus *2NLC > NLC + LM*(*c_in_ + c_out_*) ⇔ *NC > M*(*c_in_ + c_out_*) which is true for circuits with small *M, c_in_, c_out_*.

2 The tensor weights are very high-dimensional so that the least-squares method and variations thereof have failed due to the high memory requirements.

## Notes

### Competing Interest Statement

The authors have declared no competing interest.

