## Supplementary Figures for "Single circuit in V1 capable of switching contexts during movement using VIP population as a switch"

### 1 Supplemental figures

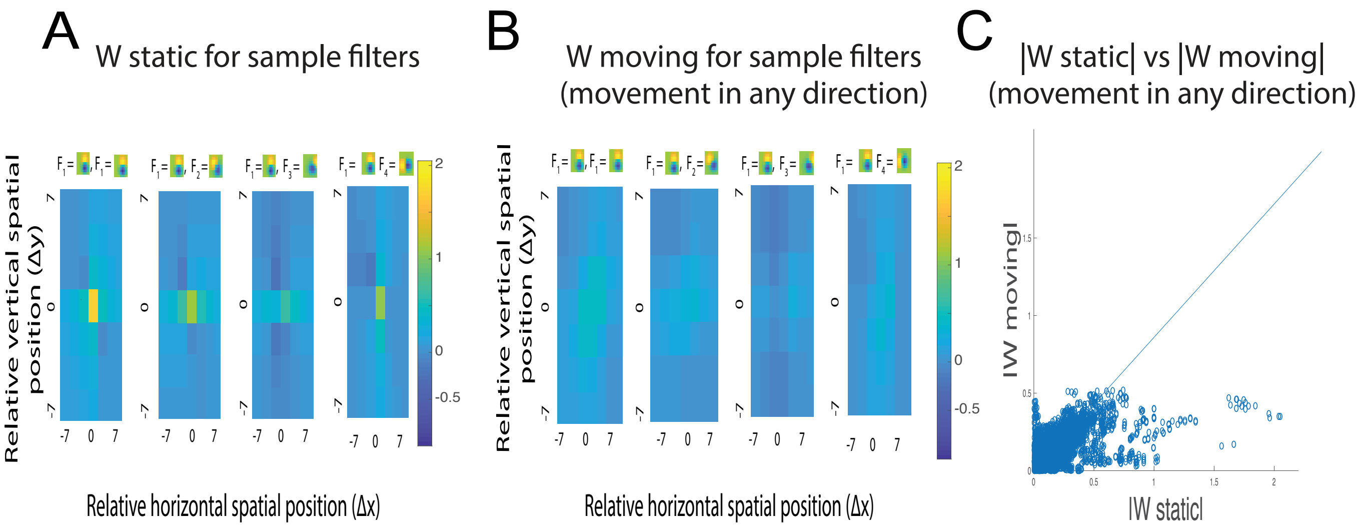


**Figure S1:** a. Slices of $\mathbf{W}^{\text{static}}$ corresponding to different pairs of filters (feature $\mathbf{F}_1$ paired with features $\mathbf{F}_1$ - $\mathbf{F}_4$). b. Slices of $\mathbf{W}^{\text{moving}}$ computed for dataset of videos where movement is in any direction. Slices shown correspond to different pairs of filters (feature $\mathbf{F}_1$ paired with features $\mathbf{F}_1$ - $\mathbf{F}_4$). c. Scatter plot of $|\mathbf{W}^{\text{static}}|$ vs $|\mathbf{W}^{\text{moving}}|$. This reveals that on average, $\|\mathbf{W}^{\text{static}}\| > \|\mathbf{W}^{\text{moving}}\|$ for this dataset of natural images and videos where movement can be in any direction.

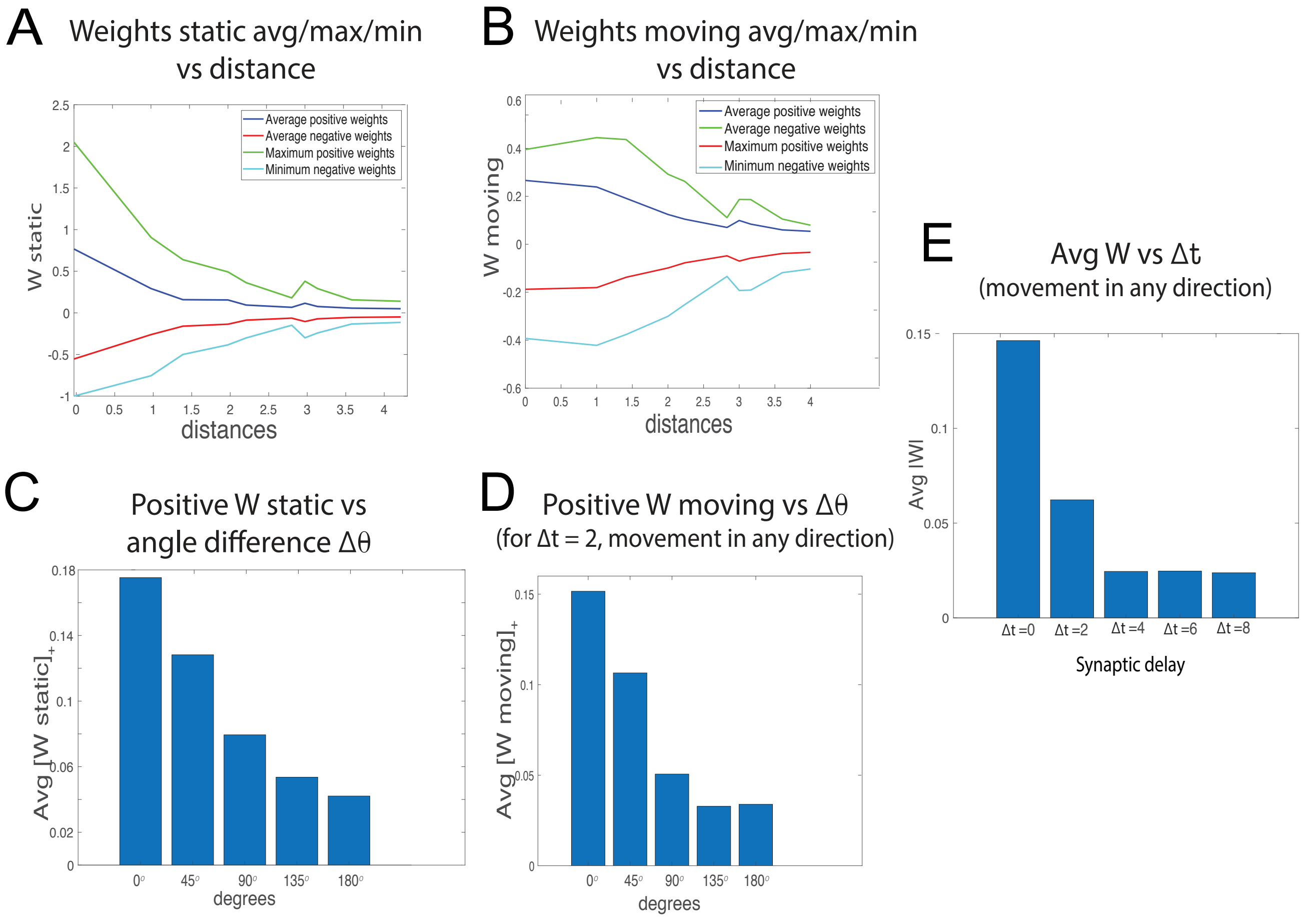


**Figure S2:** a. Dependence of the maximum, minimum, average positive and negative synaptic weights for the *static* context onto a target neuron $k$ from all neurons on the distance measured in terms of receptive field size (1 unit $= 1/2$ RF size $= 7$ pixels). This distance dependence enables us to compute the spatial constant in terms of the classical receptive field size and compare it to data. b. Dependence of the maximum, minimum, average positive and negative synaptic weights for the *moving* context ($\Delta t = 2$) onto a target neuron $k$ from all neurons on the distance measured in terms of receptive field size (1 unit $= 1/2$ RF size $= 7$ pixels). The dataset of videos used to compute the weights here and in d, e is the one where the movement can be in any direction. c. Predicted average positive synaptic weight in the static context as a function of difference in orientation of features. This predicts that excitatory weights between neurons responsive to more similar features (similar in orientation) are stronger than those between neurons responsive to different features. The trend matches data in [1]. d. Predicted average positive synaptic weight in the moving context ($\Delta t = 2$) as a function of difference in orientation of features. e. Average strength of moving synaptic weights as a function of $\Delta t$, a parameter describing synaptic delay. The higher the synaptic delay, the closer to chance the co-occurrence probability is, and thus the lower the absolute values of the synaptic weights are.

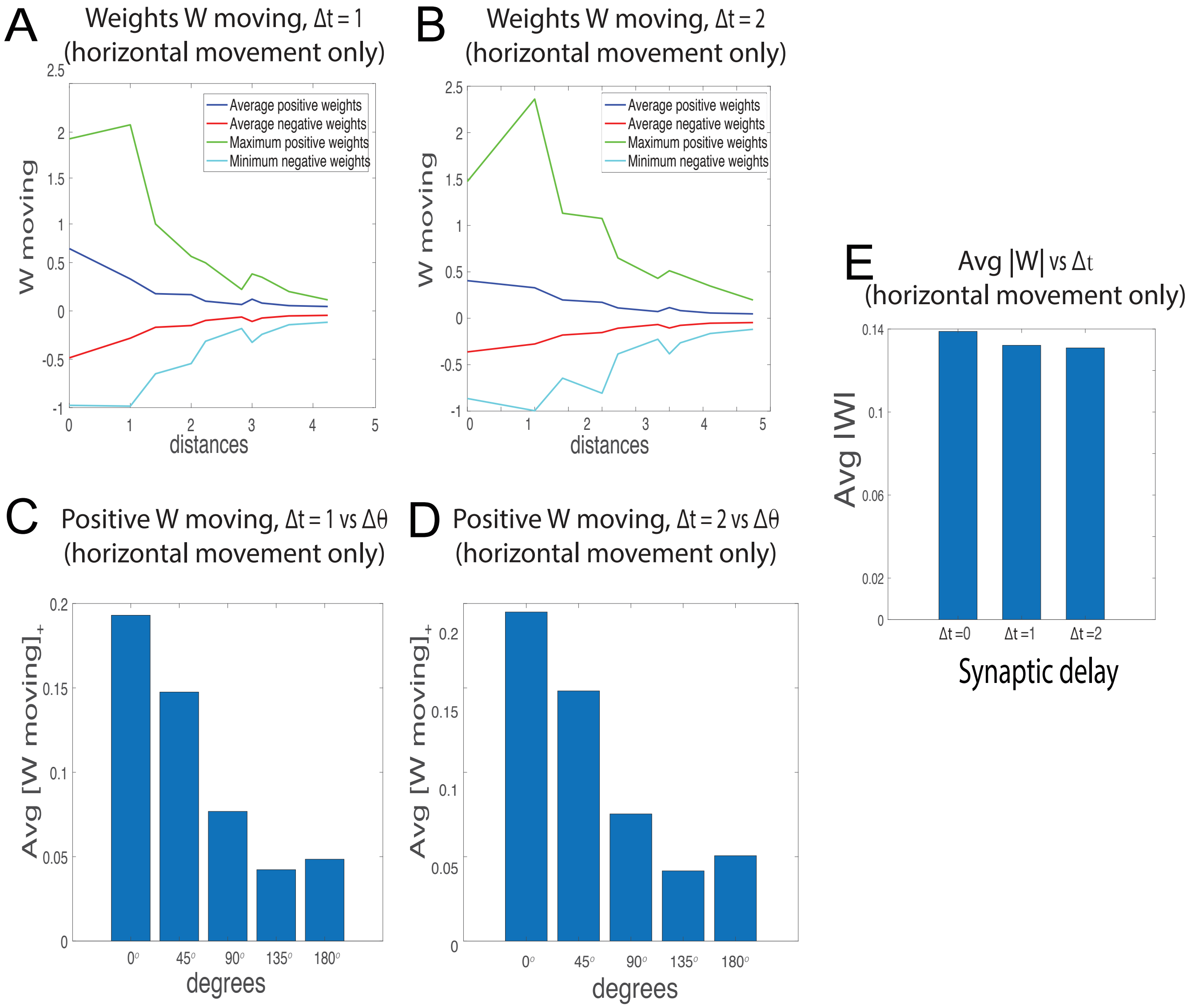


**Figure S3:** a. Dependence of the maximum, minimum, average positive and negative synaptic weights for the *moving* context with $\Delta t = 1$ onto a target neuron $k$ from all neurons on the distance measured in terms of receptive field size (1 unit = 1/2 RF size = 7 pixels). The dataset of videos used to compute the weights here and throughout this figure is the one where the movement can be only in the horizontal rightward direction. Because the movement is 3 pixels/frame and $\Delta t = 1$ frame, the peak weight is between neurons responding preferentially to identical features and classical receptive fields centered 3 pixels apart (i.e. peak is at $\mathbf{W}^{3,0}_{kk}$, where $\Delta x = 3 \approx 1/4$ RF $= 1/2$ unit distance, not shown in the plot). b. Dependence of the maximum, minimum, average positive and negative synaptic weights for the *moving* context with $\Delta t = 2$ onto a target neuron $k$ from all neurons on the distance measured in terms of receptive field size (1 unit = 1/2 RF size = 7 pixels). Because the movement is 3 pixels/frame and $\Delta t = 2$ frames, the peak weight is between neurons responding preferentially to identical features and classical receptive fields centered 6 pixels apart (i.e. peak is at $\mathbf{W}^{6,0}_{kk}$, where $\Delta x = 6 \approx 1/2$ RF $= 1$ unit distance). c. Predicted average positive synaptic weight in the moving context ($\Delta t = 1$) as a function of difference in orientation of features. d. Same as c, but with $\Delta t = 2$. e. Average weight strength in terms of synaptic delay $\Delta t$, where $\Delta t = 0$ corresponds to $\mathbf{W}^{\text{static}}$. Unlike the weights in Figure Fig. S2, which correspond to movement in any direction, the average weight strength does not decrease significantly with $\Delta t$. Indeed, the peak of the tensor simply shifts at different spatial positions depending on how large the synaptic delay is, but otherwise the tensor remains (mostly) unchanged.

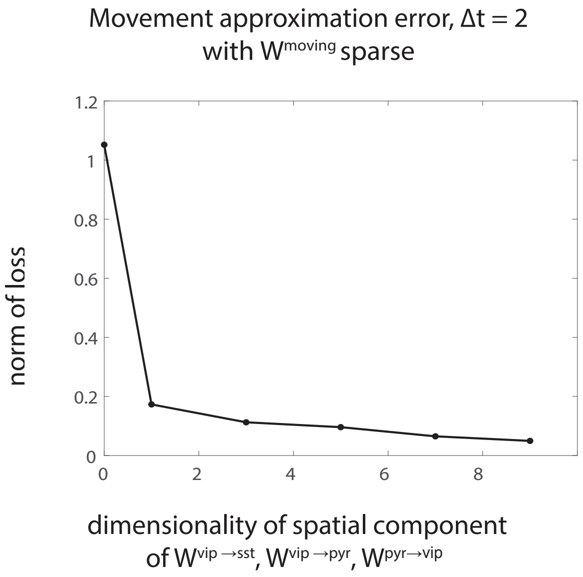


**Figure S4:** Varying the dimensionality of the tensors $\mathbf{W}^{\mathrm{VIP}\to\mathrm{SST}}$, $\mathbf{W}^{\mathrm{VIP}\to\mathrm{PYR}}$, $\mathbf{W}^{\mathrm{PYR}\to\mathrm{VIP}}$ can lower the movement approximation error as defined in (**??**). These tensors have dimension $Nf_1 \times Nf_2 \times c \times c$, where $Nf_1, Nf_2$ represent the number of VIP, SST, or PYR neurons, and $c$ represents the dimensionality corresponding to the spatial component (shown on x-axis). We set $\Delta t = 2$, $Nf_1 = 5, Nf_2 = 34$ for $\mathbf{W}^{\mathrm{VIP}\to\mathrm{SST}}$, $\mathbf{W}^{\mathrm{VIP}\to\mathrm{PYR}}$, $Nf_1 = 34, Nf_2 = 5$ for $\mathbf{W}^{\mathrm{PYR}\to\mathrm{VIP}}$, and use sparse weights for the optimization procedure.

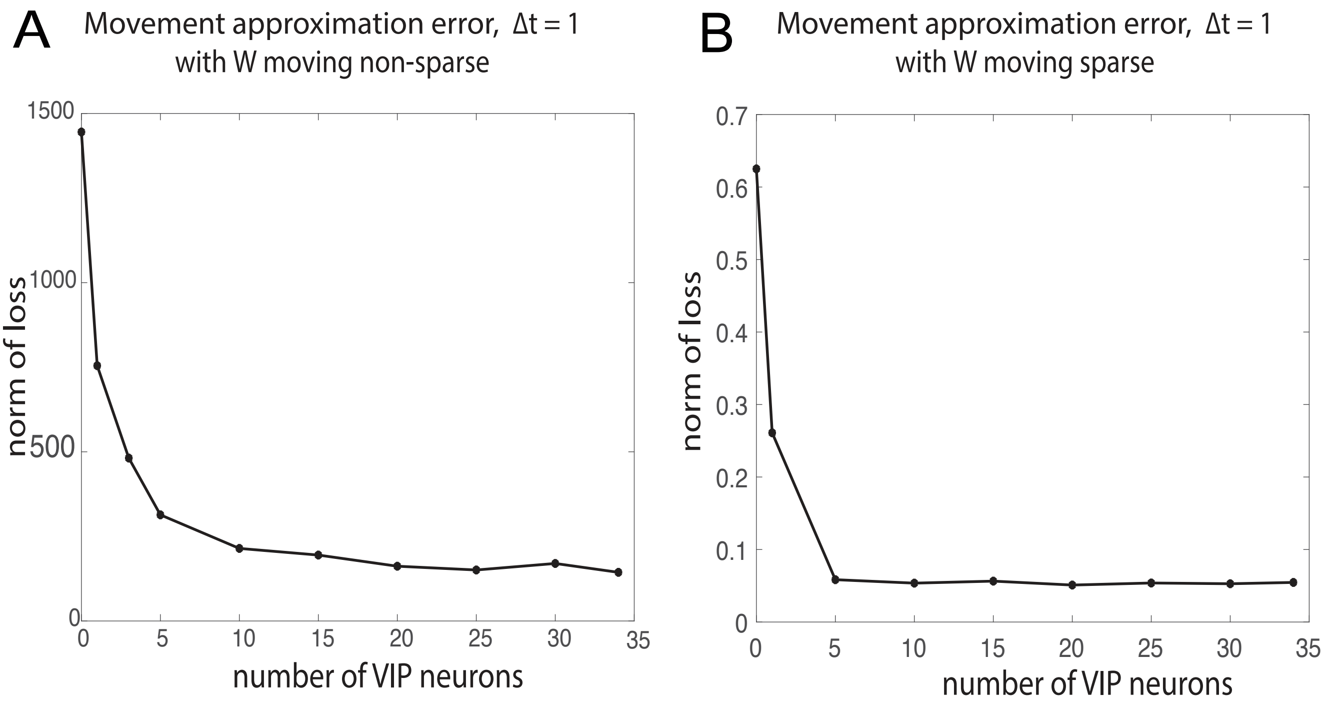


**Figure S5:** a. Movement approximation error (defined as in (**??**)) decreases with increasing number of VIP neurons for synaptic delay $\Delta t = 1$ and using the full $\mathbf{W}^{\text{moving}}$ (non-sparse). b. Movement approximation error decreases with increasing number of VIP neurons for synaptic delay $\Delta t = 1$ and using the sparse sampled $\mathbf{W}^{\text{moving}}$.

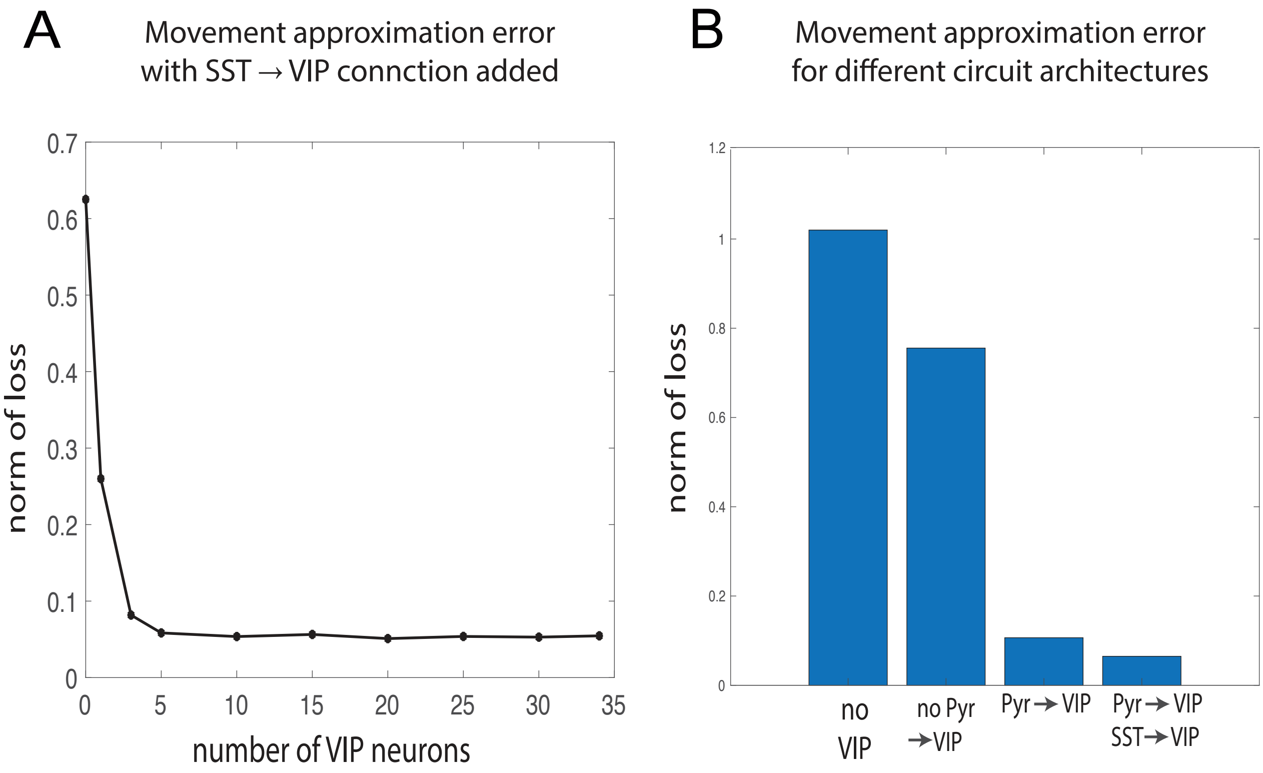


**Figure S6:** a. Movement approximation error (defined as in (**??**)) decreases with increasing number of VIP neurons, after an additional connection from SST to VIP is added. We set synaptic delay to $\Delta t = 1$ and use the sparse sampled $\mathbf{W}^{\text{moving}}$. b. Movement approximation error for different circuits: a circuit with no VIP units (leftmost bar), a circuit with VIP and connections from VIP to PYR and SST (middle left bar), a circuit with an additional connection from PYR to VIP added (middle right bar), a circuit with an additional connection from SST to VIP added (rightmost bar).

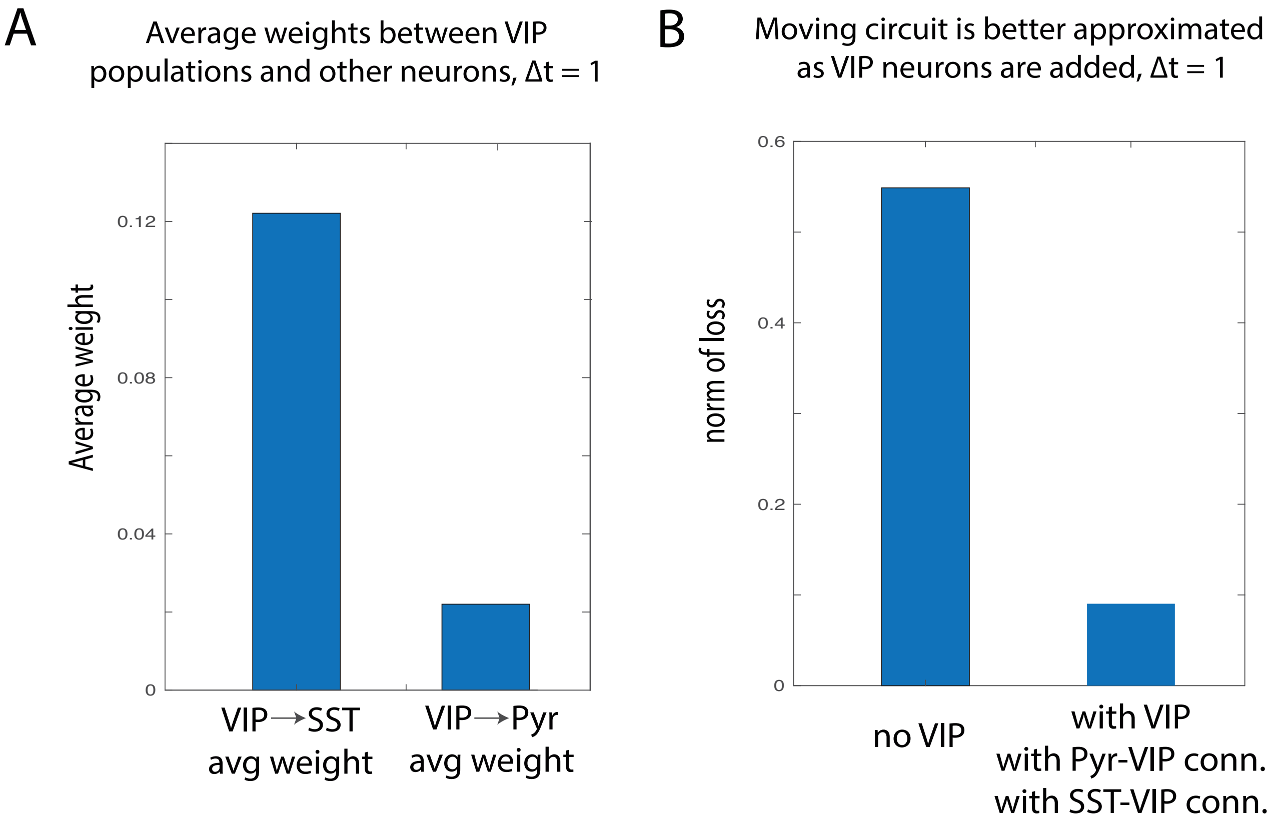


**Figure S7:** a. Comparison of $\mathbf{W}^{VIP\rightarrow SST}$ average weights to $\mathbf{W}^{VIP\rightarrow PYR}$ average weights (0.12 compared to 0.022). The ratio between these average weights is invariant to re-scaling due to patch independence that results in sparse weights $\mathbf{W}^{VIP\rightarrow SST}, \mathbf{W}^{VIP\rightarrow PYR}$. These weights have been computed by optimizing (**??**) for $\mathbf{W}^{\text{moving}}$ with $\Delta t = 1$ (although a similar result holds for $\Delta t = 2$) b. Verifying that using the solutions $\mathbf{W}^{VIP\rightarrow SST}, \mathbf{W}^{VIP\rightarrow PYR}$ to the optimization problem (**??**) yields a small movement approximation error (right bar) compared to the same error $E_{switch,2}$ when no VIP units are considered (left bar). The movement approximation error when VIP units are added (right bar) is for the circuit that includes SST to VIP and PYR to VIP connections.

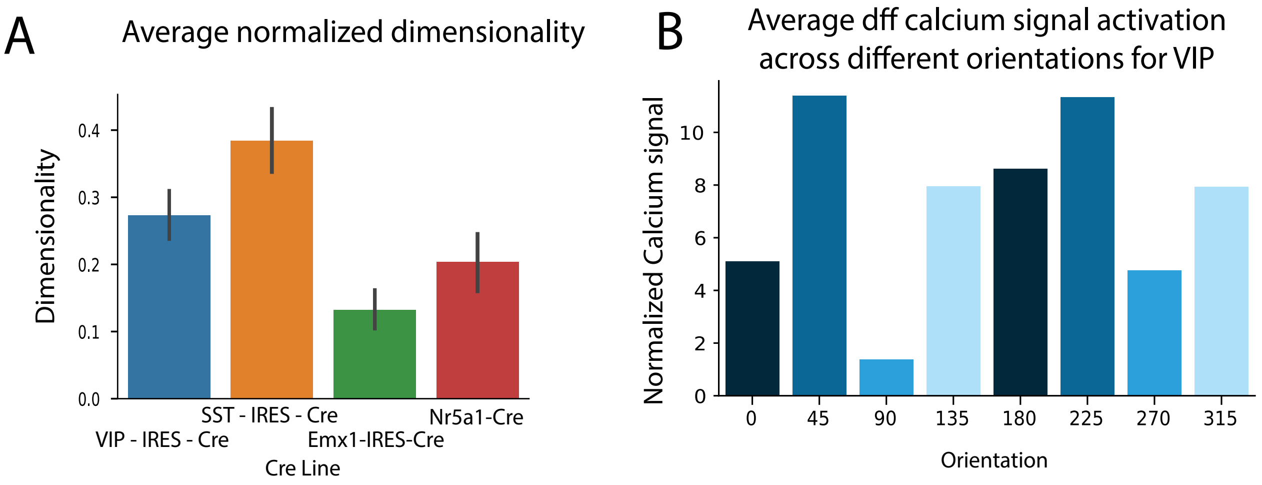


**Figure S8:** a. Average dimensionality across sessions normalized to the number of neurons in each session for multiple neural populations. Dimensionality is assessed by means of the measure Participation Ratio during epochs of spontaneous activity for the dff signal of calcium. While the average dimensionality of the activity of the PYR population is lower, this is partially due to the number of PYR units recorded being higher. b. Average dff calcium signal activation across different orientations for the VIP population during drifting gratings stimuli. Despite the trend appearing across orientations this is not significant as the Standard Error (not shown) is high due to the high variability across recordings.
